# Comparing the transmission of carbapenemase-producing and extended-spectrum beta-lactamase-producing *Escherichia coli* between broiler chickens

**DOI:** 10.1101/2023.02.21.529369

**Authors:** Natcha Dankittipong, Jesse B. Alderliesten, Jan Van den Broek, M. Anita Dame-Korevaar, Michael S. M. Brouwer, Francisca C. Velkers, Alex Bossers, Clazien J. de Vos, Jaap A. Wagenaar, J. Arjan Stegeman, Egil A. J. Fischer

**Author notes:** Biosystems Data Analysis, Swammerdam Institute for Life Sciences, University of Amsterdam, Sciencepark 904, Amsterdam, The Netherlands (Jesse B. Alderliesten). These authors contributed equally (Natcha Dankittipong, Jesse B. Alderliesten).

## Abstract

The emergence of carbapenemase-producing *Enterobacteriaceae* (CPE) is a threat to public health, because of their resistance to clinically important carbapenem antibiotics. The emergence of CPE in meat-producing animals is particularly worrying because consumption of meat contaminated with resistant bacteria similar to CPE, such as extended-spectrum beta-lactamase (ESBL)-producing Enterobacteriaceae, contributed to colonization in humans worldwide. Currently, no data on the transmission of CPE in livestock is available. We performed a transmission experiment to quantify the transmission of CPE between broilers to fill this knowledge gap and to compare the transmission rates of CPE and other antibiotic-resistant *E. coli*.

A total of 180 Ross 308 broiler chickens were distributed on the day of hatch (day 0) over 12 pens. On day 5, half of the chickens in each pen were orally inoculated with 5·10^2^ colony-forming units of CPE, ESBL, or chloramphenicol-resistant *E. coli* (catA1). Amoxicillin drinking water treatment was given twice daily in 6 of the 12 pens from days 2 to 6 to evaluate the effect of antibiotic treatment on the transmission rates. Cloacal swabs of all animals were taken to determine the number of infectious broilers. We used a Bayesian hierarchical model to quantify the transmission of the *E. coli* strains. *E. coli* can survive in the environment and serve as a reservoir. Therefore, the susceptible-infectious transmission model was adapted to account for the transmission of resistant bacteria from the environment. In addition, the caecal microbiome was analyzed on day 5 and at the end of the experiment on day 14 to assess the relationship between the caecal microbiome and the transmission rates.

The transmission rates of CPE were 52 – 68 per cent lower compared to ESBL and catA1, but it is not clear if these differences were caused by differences between the resistance genes or between the *E. coli* strains. Differences between the groups in transmission rates and microbiome diversity did not correspond to each other, indicating that differences in transmission rates were probably not caused by major differences in the community structure in the caecal microbiome. Amoxicillin treatment from day 2 to 6 increased the transmission rate more than three-fold in all inoculums. It also increased alpha-diversity compared to untreated animals on day 5, but not on day 14, suggesting only a temporary effect.

Future research could incorporate more complex transmission models with different species of resistant bacteria into the Bayesian hierarchical model.

## 1. Introduction

Carbapenemase-producing *Enterobacteriaceae* (CPE; also referred to as carbapenem-resistant *Enterobacteriaceae*) are potentially life-threatening bacteria because of their resistance to clinically important carbapenem antibiotics (Brink, 2019; World Health Organization, 2019; Zhou et al., 2021). CPE are detected worldwide in farm animals, wild animals, companion animals, fish, and the environment (Köck et al., 2018; Bonardi and Pitino, 2019). The emergence of CPE in meat-producing animals is particularly worrying because consumption of meat contaminated with resistant bacteria similar to CPE, such as extended-spectrum beta-lactamase (ESBL)-producing bacteria or plasmid-encoded AmpC (pAmpC)-producing bacteria, contributes to colonization in humans worldwide (Leverstein-van Hall et al., 2011; Rousham et al., 2018; Mughini-Gras et al., 2019). Consequently, it is crucial to assess the transmission dynamics of CPE in livestock farms. We looked at transmission between broilers because the prevalence of ESBL-producing bacteria in broilers is high compared to other livestock (European Food Safety Authority and European Centre for Disease Prevention Control, 2022).

The transmission rate parameter *β* is a key parameter to describe the transmission dynamics in populations and is here defined as the rate of successful transmission per time unit following contact with an infectious source such as bacteria carrying resistance genes (Keeling and Rohani, 2007). Transmission of ESBL-producing *Escherichia coli (E. coli)* in poultry has been investigated extensively (Huijbers et al., 2016; Dame-Korevaar et al., 2019; Robé et al., 2019; Dame-Korevaar et al., 2020a; Dame-Korevaar et al., 2020b), showing among others that 2 strains of beta-lactamase-producing bacteria (carrying *bla*_CTX-M-1_ and *bla*_CMY-2_, respectively) colonized broilers at the same rate (Dame-Korevaar et al., 2019). In contrast, no data on the transmission of CPE in livestock is available. Although poultry is at risk of CPE introduction (Dankittipong et al., 2022), the prevalence of CPE in animals is much lower than the prevalence of ESBL/pAmpC-producing bacteria (European Centre for Disease Prevention and Control, 2018). The difference in the prevalence of CPE and ESBL-producing bacteria could be explained by differences in the transmission dynamics of the resistance genes and the plasmids that carry them (Rozwandowicz et al., 2018; Wilson and Török, 2018). Differences in selective pressure caused by historical use in livestock of third-generation cephalosporins that co-select for carbapenemase-producing genes (Ogunrinu et al., 2020) compared to the use of carbapenems having worldwide never been allowed in livestock (Madec and Haenni, 2018) might also contribute to the difference in prevalence.

Conventional methods to quantify the transmission of bacteria assume direct transmission between animals (Velthuis et al., 2007). However, *E. coli* can survive for a considerable amount of time in the environment (Table S14) and is commonly transmitted between animals through the faecal-oral route (Lister and Barrow, 2008; van Elsas et al., 2011; van Bunnik et al., 2014). Previous transmission experiments of ESBL-producing bacteria in broilers, wildtype nalidixic-resistant *E. coli* in broilers, and *Salmonella Dublin* in young dairy calves highlighted the excretion of these bacteria into the environment and subsequent acquisition of excreted bacteria from the environment as a key mechanism of transmission (Nielsen et al., 2007; van Bunnik et al., 2014; Dame-Korevaar et al., 2017).

Antibiotic usage is a primary driver of resistant bacteria in clinical and non-clinical settings (Knobler et al., 2003; Davies and Davies, 2010; Holmes et al., 2016) and is widespread in livestock worldwide (Mathew et al., 2007; Aarestrup, 2015). Twenty-two per cent of the conventional broiler farms in the Netherlands did not use antibiotics in 2020, but 44% had a persistently high antibiotic usage exceeding the action threshold defined by the Netherlands Veterinary Medicines Institute and 5% had a persistently high antibiotic usage exceeding the sector-negotiated action threshold (Bonten et al., 2021). Treatment with antibiotics generally temporarily decreases the number of bacterial species in the gut microbiome and lowers the abundance of some common taxa, allowing the abundance of some low-abundant taxa or opportunistic pathogens to increase (Kim et al., 2017; Rochegüe et al., 2021). This might affect the transmission of bacteria, because a more diverse gut microbiome hinders colonization by exogenous bacteria (Kim et al., 2017; Sorbara and Pamer, 2019), thereby reducing the excretion of these bacteria (Dame-Korevaar et al., 2020b).

We performed a transmission experiment to quantify the transmission of CPE between broilers and to quantitatively compare the transmission rates of CPE and ESBL-producing *E. coli* to determine if the difference in the prevalence of CPE and ESBL in broilers might have been caused by differences in the transmission dynamics of the resistance genes and the plasmids that carry them. Groups with and without amoxicillin treatment were compared to investigate if and how antibiotic treatment affects the transmission, and relations between differences in transmission rates and the caecal microbiome were assessed.

## 2. Material and Method

### 2.1. Transmission experiment

The study protocol was approved by the Dutch Central Authority for Scientific Procedures on Animals and the Animal Experiments Committee of Utrecht University (Utrecht, the Netherlands) under registration number AVD108002015314 and all procedures were performed in full compliance with all legislation. All broilers were observed daily, and any abnormality and mortality were recorded.

#### 2.1.1. Inoculums

Three inoculums were prepared for this experiment, referred to as the CPE-strain, ESBL-strain, and catA1-strain throughout the paper (Table 1). All strains were *E. coli* obtained from broilers in conventional farms in Europe. Before inoculation, all strains were streaked on heart infusion agar with 5% sheep blood (Becton Dickinson GmbH, Heidelberg, Germany), transferred to LB medium, and cultured overnight. The *E. coli* cultures were diluted in phosphate-buffered saline with 0.5 McFarland standards resulting in 1.10 · 10^8^ bacteria suspension per mL. Prepared inoculums were enumerated in duplicate counts and each contained 0.55 · 10^3^ – 1.0 · 10^3^ colony-forming units per mL.

**Table 1:** Characteristics of the CPE, ESBL, and catA1 isolates used as inoculums. Abbreviations: Inc-group: incompatibility group; MLST: multi-locus sequence type.

| Inoculum | *E. coli* isolate | MLST | Selected resistance | Gene | Plasmid Inc-group | Host's country of origin | Reference |
| --- | --- | --- | --- | --- | --- | --- | --- |
| CPE-strain | CFSAN083827 | 4980 | Carbapenem | OXA-162 | HI2 | Romania | (Bortolaia et al., 2021) |
| ESBL-strain | SafeFoodEra-230 | 101 | Extended-spectrum beta-lactam | CTX-M2 | HI2 | Germany | (Wu et al., 2013) |
| catA1-strain | EFFORT 102803008 | 10 | Chloramphenicol | catA1 | FIB/FII | The Netherlands | (Leekitcharoenphon et al., 2021) |

#### 2.1.2. Sampling scheme and experimental design

The experiment was conducted in human Biosafety level 3 (BSL-3) facilities at Wageningen Bioveterinary Research (WBVR), Lelystad. Before the experiment, samples from the parent stock and environmental samples from the incubator (BSL-1) and experimental facilities were taken which confirmed the absence of ESBL-producing *E. coli*. Two hundred and forty eggs were collected from a conventional Ross 308 broiler parent stock, individually disinfected with 3% hydrogen peroxide and incubated for 21 days at BSL-1 experimental facilities of WBVR. On the day of hatch, day 0, 180 hatchlings were transported to the BSL-3 animal facilities of WBVR, where they were weighted, neck tagged with an individual number and randomly distributed over 12 pens, with 15 unsexed broilers per pen (see Table S3 for an overview of the distribution of the sexes in the different groups). Broilers of both sexes were used because a mixed group reflects the practical situation in terms of group dynamics and the prevalence of ESBL or CPE is not known to differ by gender. Pens had a surface area of 1.35 m^2^, with a bedding of sterilized wood shavings, and were separated from each other by fences of 70 – 80 cm high such that no direct contact was possible between pens. Broilers had *ad libitum* access to feed and water and a standard lighting and temperature scheme for broiler chickens was used. The feed should have been a standard broiler diet without antibiotics or coccidiostats, but accidentally feed for layer pullets, free of antibiotics and coccidiostats, was provided. The feed was based on wheat, maize, and soybean meal and contained 2,563 kcal of apparent metabolizable energy per kg and 20% of crude protein heated to 90 °C. From days 2 to 6, amoxicillin was provided via drinking water twice a day at the suppliers’ recommended dose of 20 mg/kg live weight to the broilers in pens 3, 4, 7, 8, 11, and 12 (Figure 1). Amoxicillin was used as an example of a broad-spectrum antibiotic commonly used in broilers (Ventola, 2015) to compare the transmission of all inoculums in the absence and presence of antibiotic treatment.

**Figure 1:**
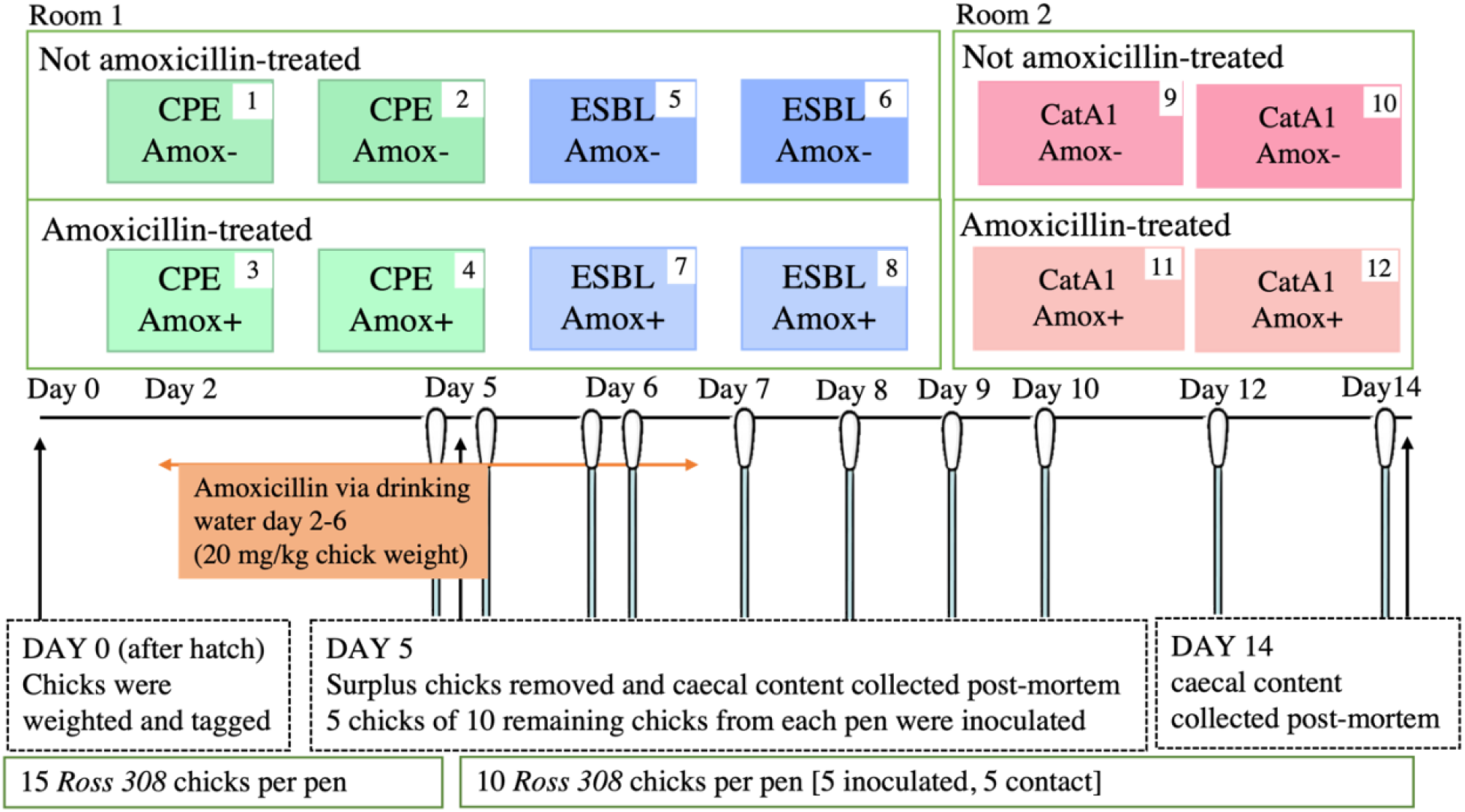
Setup of the pens (top) and timeline of the experimental design from the moment of hatch to the end of the experiment on day 14, with the sampling time points indicated by the swabs (bottom). Abbreviations: Amox-: non-amoxicillin-treated; Amox+: amoxicillin-treated.

On day 5, cloacal swabs were taken from all broilers using sterile dry Eswabs (MW100, Medical Wire & Equipment, England) to confirm the absence of CPE and ESBL-producing *E. coli*. 10 broilers per pen were kept for the transmission experiment and surplus broilers (at most 5 per pen) were euthanized and their caecal content was collected for microbiome analysis. Five broilers randomly chosen out of the 10 remaining broilers per pen were separated from the other broilers and orally inoculated (using a syringe with a crop needle) with 0.5 mL PBS which per mL containing approximately 10^3^ colony-forming units of *E. coli*, i.e., the CPE-strain (pens 1 – 4), the ESBL-strain (pens 5 – 8), or the catA1-strain (pens 9 – 12). One hour after inoculation, inoculated broilers were returned to their pen where they resided with contact broilers (i.e., broilers that were not inoculated). Cloacal swabs were taken from all broilers at approximately 8 hours after inoculation on day 5, twice on day 6 (8 hours apart), and once per day on days 7 to 10, 12, and 14 (Figure 1) (Dame-Korevaar et al., 2020a). All broilers were euthanized on day 14 and their caecal content was collected for microbiome analysis.

#### 2.1.3. Resistance gene detection

All cloacal swabs were enriched overnight in 3 mL buffered peptone water at 37 °C. Thereafter they were inoculated onto selective MacConkey plates supplemented with 0.5 mg/L ertapenem (swabs from pens 1 – 4), 1 mg/L cefotaxime (swabs from pens 5 – 8), or 64 mg/L chloramphenicol (swabs from pens 9 – 12) using a sterile loop and incubated overnight at 37 °C. A broiler was defined as positive when colonies were detected on MacConkey plates after overnight incubation. The pen, used inoculum, antibiotic treatment, and the test results of the cloacal swabs (i.e., positive or negative for CPE-strain, ESBL-strain, or catA1-strain) at each sampling time point were recorded for all inoculated and contact broilers (Table S1).

#### 2.1.4. Microbiome sequencing

Microbial DNA was isolated from 0.2 g caecal content according to the manufacturer’s instructions using the PureLink microbial DNA isolation kit (Thermo Fisher Scientific, Waltham, Massachusetts, USA). Negative controls spiked with a low concentration of microbial community DNA standard (ZymoBIOMICS; Zymo Research Corporation, Irvine, CA) were used in the batches of DNA isolation and amplification thereafter as control of performance and sanity throughout the processing (see Figure S1 for a comparison of the theoretical and obtained composition of the negative controls). Following extraction, the DNA extracts were quantified with an InvitrogenTM QubitTM 3.0 Fluorometer and stored at -20 °C for further processing. The hypervariable regions V3+V4 of the 16S rRNA gene were amplified in triplicate using a limited-cycles PCR with the primers CVI_V3-forw CCTACGGGAGGCAGCAG and CVI_V4-rev GGACTACHVGGGTWTCT. The following amplification conditions were used as previously described (Jurburg et al., 2019): 98 °C for 2 minutes, followed by 20 cycles of 98 °C for 10 s, 55 °C for 30 s, and 72 °C for 10 s, and finally by 72 °C for 7 minutes. Triplicate PCR products were pooled per sample and checked on a TapeStation (Agilent, USA) and after barcode indexing subsequently sequenced on a MiSeq sequencer (Illumina Inc., San Diego, CA) using a version 3 paired-end 300 bp kit.

### 2.2. Data analysis

#### 2.2.1. SI-model

The transmission of *E. coli* between broilers was modelled using a compartmental susceptible-infectious model (SI-model; Figure 2). Previous research identified excretion and subsequent acquisition of *E. coli* from the environment as a key mechanism of transmission (Lister and Barrow, 2008; van Bunnik et al., 2014). We incorporated this in our SI-model by assuming excreting broilers (I) excrete viable bacteria into the environment of their pen at a constant rate of ω units per hour from the moment they start to excrete, and these excreted bacteria will decay at a rate *δ*. The unknown excretion rate (ω) was scaled such that the hazard produced by 1 broiler during 1 time unit is 1 (Gerhards et al., 2022). The environmental hazard at time t is denoted as *E*_*t*_. A detailed description of the model including the scaling is given in Section 3.1 of the supplementary information.

**Figure 2:**
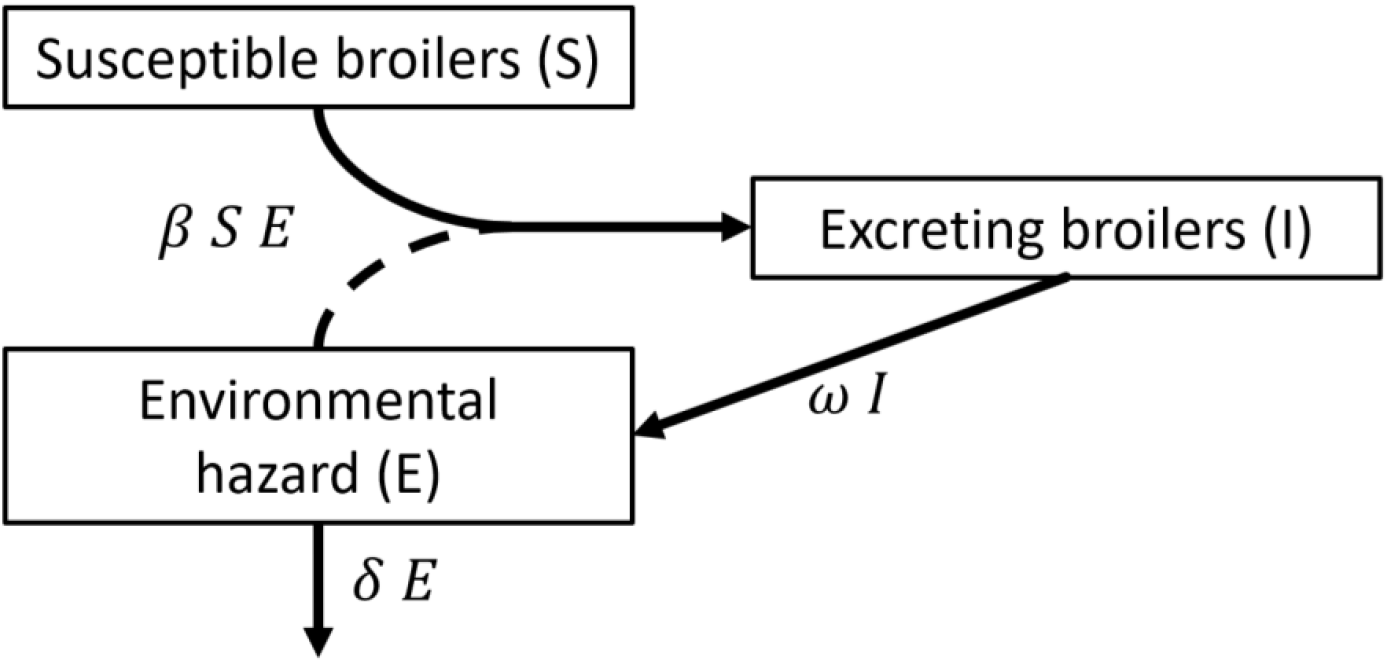
compartmental SI-model of indirect transmission of E. coli between broilers. Excreting broilers (I; positive inoculated broilers and positive contact broilers) excrete bacteria into the environment at rate ω. Only negative contact broilers are counted as susceptible broilers (S) because negative inoculated broilers are assumed to start excreting through inoculation instead of through colonization after contact with the environmental hazard (E). Environmental hazard decays at rate δ (h^-1^). Susceptible contact broilers become colonized through contact with bacteria in the environment at transmission rate parameter β (h^-1^), thus becoming excreting broilers. The dashed line connecting environmental hazard and excreting broilers indicates bacteria in the environment facilitate colonization but are not converted to excreting broilers.

When negative contact broilers were colonized through contact with bacteria in the environment at rate *β S*_*t*_ *E*_*t*_, they were denoted as cases and incorporated in the SI-model as excreting for the next time interval. Negative inoculated broilers were assumed to start excreting through inoculation instead of through contact with bacteria in the environment and were therefore not denoted as cases.

In the SI-model it is assumed that contact broilers are either susceptible (S) or excreting (I). Once broilers start excreting, it is assumed they will continue to excrete until the end of the experiment. To adhere to this structure, a negative test result in a broiler that previously tested positive was assumed to be false negative (see section 1.2 with Table S2 in the supplementary material). In pens 3, 4, 11 and 12, the first positive tests for inoculated and contact broilers occurred at the same time point. However, at least one inoculated broiler must start excreting before colonization of contact broilers can occur. Therefore, we assumed inoculated broilers started excreting halfway between the first time point they tested positive and the previous sampling time point, and contact broilers were assumed to start excreting slightly slower, from the time point they tested positive.

#### 2.2.2. Bayesian hierarchical inference

A Bayesian hierarchical model was used to infer the parameters of the SI-model (see section 3.2 in the supplementary material), which requires prior probability distributions for the parameters, observed data (i.e., the number of positive and negative broilers at each sampling time point in each pen), and a likelihood function. The transmission rate parameter (*β*), which indicates the infectivity and susceptibility of animals, was estimated for each pen separately from the number of susceptible broilers and the force of infection by estimating the average transmission rate parameter over all pens 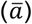 and the between-pen variation of the transmission rate parameter (*z*_*i*_). Consequently, transmission in pen i occurs at rate parameter *β*_*i*_ that is the product of the individual transmission rate parameter in that pen 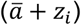 and the environmental hazard in that pen (*E*_*t*_). Posterior distributions of the transmission rate parameter for the different clusters (i.e., inoculum and antibiotic treatment) were obtained by combining the posterior distributions of 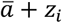 of all pens in that specific cluster.

We used results from a previous transmission study in broilers (Dame-Korevaar et al., 2020a) to define prior probability distributions (priors) for the average transmission rate parameter 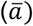 and its standard deviation (*σ*). In contrast to (Gerhards et al., 2022) we fixed the decay rate to 0.04 h^-1^ because the broilers remain excreting until the end of the experiment (Table S1) such that no information on the decay of bacteria in the environment was available.

Using the prior probabilities of the parameters and the likelihood function, parameter values were drawn using the Markov chain Monte Carlo simulated process. Four independent Markov chains (Figure S1) were initiated in the model. The transmission rate of each inoculum was extracted from the posterior distribution and transmission rates were compared using the 95% highest posterior density interval (HPDI) and the point estimate at the highest density (maximum a posteriori estimate, MAP). Differences in transmission rates between inoculums and antibiotic treatments were compared by calculating the posterior distribution of the ratio of the transmission rates.

#### 2.2.3. Microbiome analysis

The amplicon sequences were demultiplexed using *bcl2fastq* (Illumina Inc., San Diego, CA) and subsequently filtered, trimmed, error-corrected, dereplicated, chimaera-checked, and merged using R package dada2 1.16.0 (Callahan et al., 2016) with the standard parameters except for TruncLength = (270, 220), trimLeft = (25, 33), maxEE = 2 and minOverlap = 10, using a pseudo-pooling strategy. Reads were classified against the SILVA database version 138 (Quast et al., 2012). The data, the phyloseq object containing the sequence data, and the R code used for the modelling and analyses are provided at https://zenodo.org/ (DOI:xxxxxxxxx).

The number of reads in the samples (excluding negative controls) ranged from 1363 to 320392 and was standardized to 9071 reads per sample (7^th^ least number of reads; rarefy_even_depth, seed = 314; Figure S2) before alpha-diversity analysis. The final dataset contained 9540981 reads and 7952 different amplicon sequence variants (ASVs). Sequences are deposited in NCBI’s Sequence Read Archive under BioProject accession number PRJNAXXXXX.

DNA sequences isolated from caecal material obtained on days 5 and 14 were analysed separately. Non-bacterial sequences were discarded. Rarefaction curves on genus- and ASV-level were created to check if all genera and ASVs in the samples were recovered (Figure S3). Observed richness, Shannon’s index and Pielou’s evenness were used to measure alpha-diversity (Finotello et al., 2018). Kruskal-Wallis rank sum test and post hoc Dunn’s test with Benjamini-Hochberg correction were used to test for the effects of the inoculums, antibiotic treatment, and their interaction, using a significance level of 0.05. Beta-diversity, a measure of dissimilarity between communities regarding shared taxa, was analysed on non-rarefied data using Bray-Curtis dissimilarity (measuring the fraction of the community specific to either group) and Jaccard distance (measuring the fraction of taxa specific to either group, i.e., comparing presence and absence) (Schmidt et al., 2017) and visualized using the first 2 axes of the principal coordinate analysis (PCoA). Permutational multivariate analysis of variance was performed using the adonis2 function from the vegan package in R to test for effects of inoculum, antibiotic, and their interaction, and the betadisper function from the vegan package was used to test for homogeneity of group dispersions. The simper function from the vegan package was used to determine which genera contribute most to the Bray-Curtis dissimilarity between groups without and with antibiotic treatment.

#### 2.2.4. Used software

Transmission data were analysed with R version 4.1.2 (R Core Team, 2021) with package rstan 2.21.5 (Stan Development Team, 2020) using a tree depth of 14, an acceptance rate of 0.99 and 4 chains with 4000 iterations, and packages rethinking 2.21 (McElreath, 2020), cmdstanr 0.5.2 (Gabry and Cešnovar, 2022), StanHeaders 2.21.0-7 (Stan Development Team, 2018) and bayestestR 0.12.1 (Makowski et al., 2019). Sequence processing and statistical analyses related to the sequencing were performed with R 4.0.2 (R Core Team, 2020) with package dada2 1.16.0 (Callahan et al., 2016). Subsequent analyses of the microbiome data were performed with R 4.1.2 (R Core Team, 2021) with packages phyloseq 1.38.0 (McMurdie and Holmes, 2013), microbiome 1.16.0 (Lahti and Shetty, 2019), vegan 2.6.2 (Oksanen et al., 2022), and dunn.test 1.3.5 (Dinno, 2017), using packages tidyr 1.2.0 (Wickham and Girlich, 2022), dplyr 1.0.9 (Wickham et al., 2021), and Biostrings 2.62.0 (Pagès et al., 2022) for data handling, and ggplot2 3.3.6 (Wickham, 2016) and cowplot 1.1.1 (Wilke, 2020) for plotting.

## 3. Results

### 3.1. Transmission experiment

The 111 out of 120 inoculated and contact broilers that survived until the end of the experiment all became colonised by the *E. coli* strain used for inoculation (i.e., CPE-strain, ESBL-strain, or catA1-strain). Four broilers from the CPE-strain group, 4 broilers from the ESBL-strain group, and 1 broiler from the catA1-strain group died (Table S1). The majority of the broilers gained weight slower and reached 20% lower final weights than typical Ross 308 broilers, probably because they received feed for laying pullets instead of broilers. No other abnormalities were observed.

### 3.2. Transmission rates

#### 3.2.1. Predicted versus observed cases

The number of cases predicted by the hierarchical model is higher than the number of observed cases in non-antibiotic-treated pens and lower than the number of observed cases in antibiotic-treated pens because of the shrinkage caused by the hierarchical modelling (Figure 3). Shrinkage is a key feature of a hierarchical model because the measurements of different clusters (i.e., inoculum and antibiotic treatment) inform one another such that the predicted result shrinks towards the overall mean. The number of cases increased over a longer period in non-amoxicillin-treated pens (top rows) than in amoxicillin-treated pens (bottom rows) because the larger transmission rate in amoxicillin-treated pens led to the depletion of susceptible broilers.

**Figure 3:**
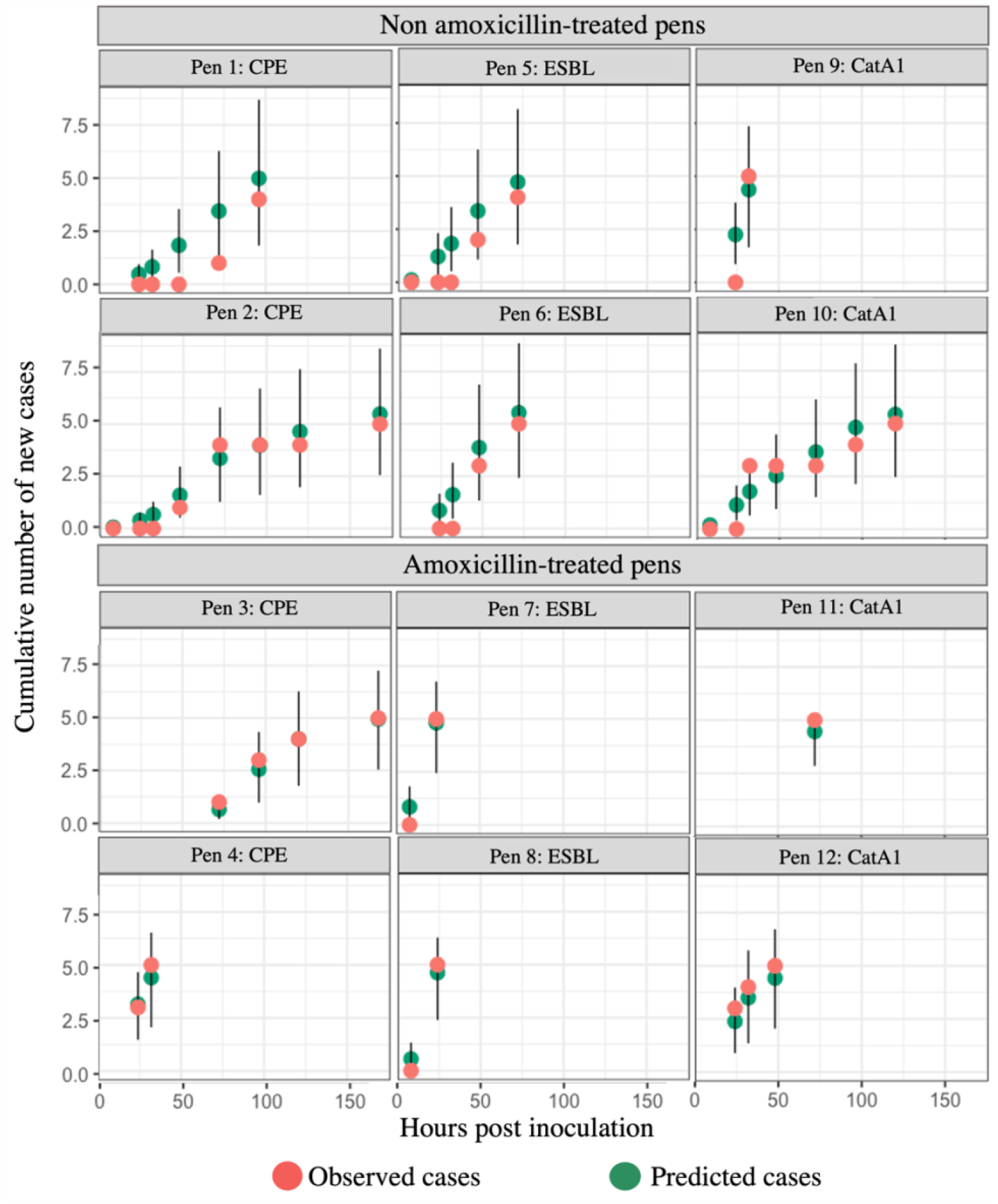
number of cases over time. Observed (orange) and predicted (green) cumulative number of new cases among the 5 susceptible broilers (i.e., susceptible contact broilers that became colonized) in each of the 12 pens (vertical axis) until the sampling time point in hours after inoculation (horizontal axis). For the predicted numbers the maximum a posteriori estimates are given, with the whiskers indicating 95% highest posterior density intervals. Transmission cannot occur when none of the broilers is excreting yet or when all broilers are excreting. No data is shown at those time points.

#### 3.2.2. Effect of inoculums

The estimated transmission rates for broilers inoculated with the CPE-strain, ESBL-strain, and catA1-strain were compared using the 95% HPDI and the MAP (shaded area and purple vertical line in Figure 4, Figure 5). In the non-amoxicillin-treated groups as well as in the amoxicillin-treated groups, the 95% HPDIs of the transmission rates of the CPE-strain, the ESBL-strain, and the catA1-strain overlap, suggesting their transmission rates are similar (Figure 4). Still, the MAP suggests that CPE-strain has the lowest transmission rate of the 3 inoculums.

**Figure 4:**
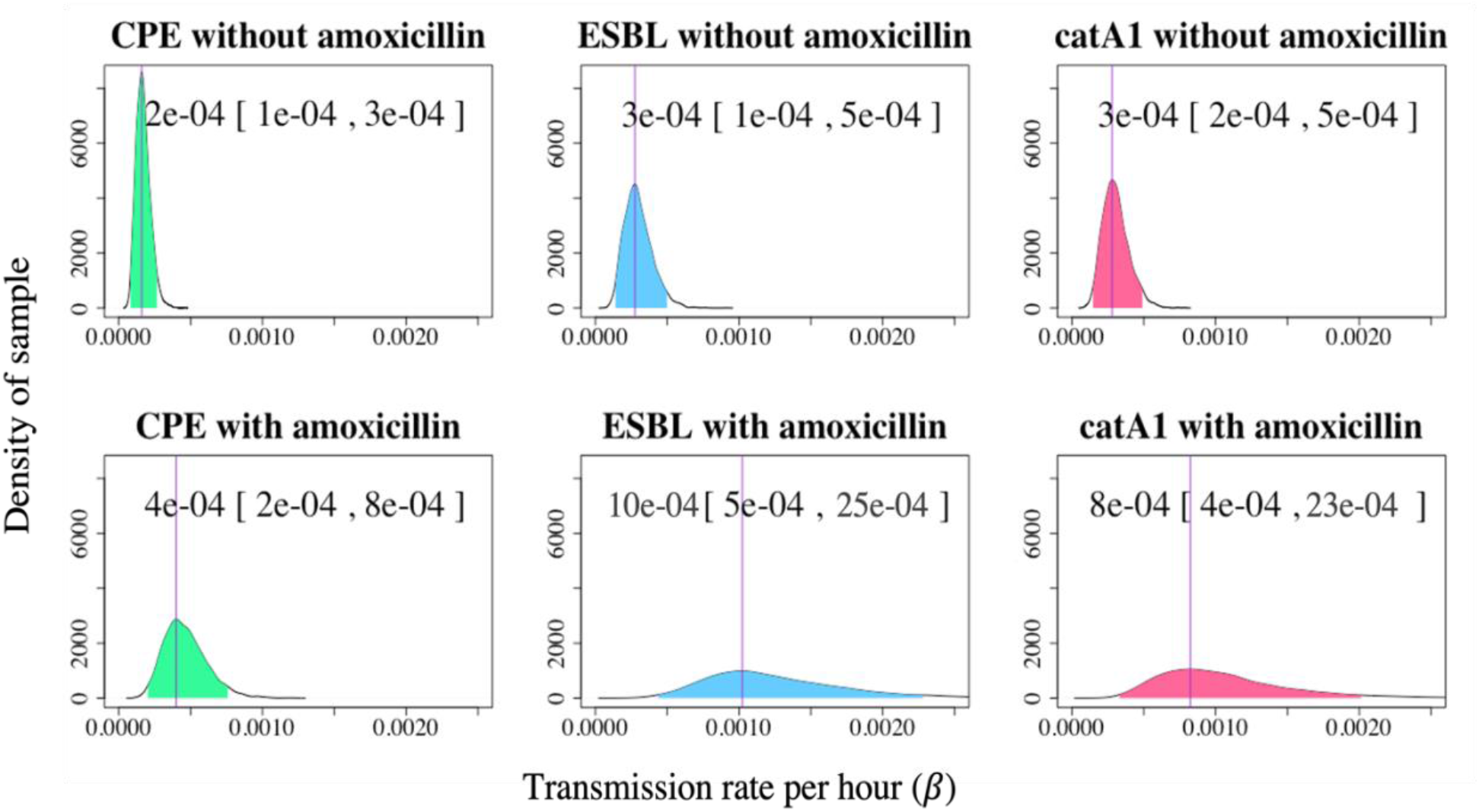
Density (vertical axis) of the posterior distribution of the transmission rate per hour (horizontal axis) for the CPE-strain, ESBL-strain and catA1-strain. The top and bottom row show plots for the pens without and with amoxicillin treatment, respectively. Purple vertical lines indicate the point estimate at the highest density and shaded areas are the 95% highest posterior density intervals of the posterior distribution; the estimated values of both are shown at the top of the plot.

**Figure 5:**
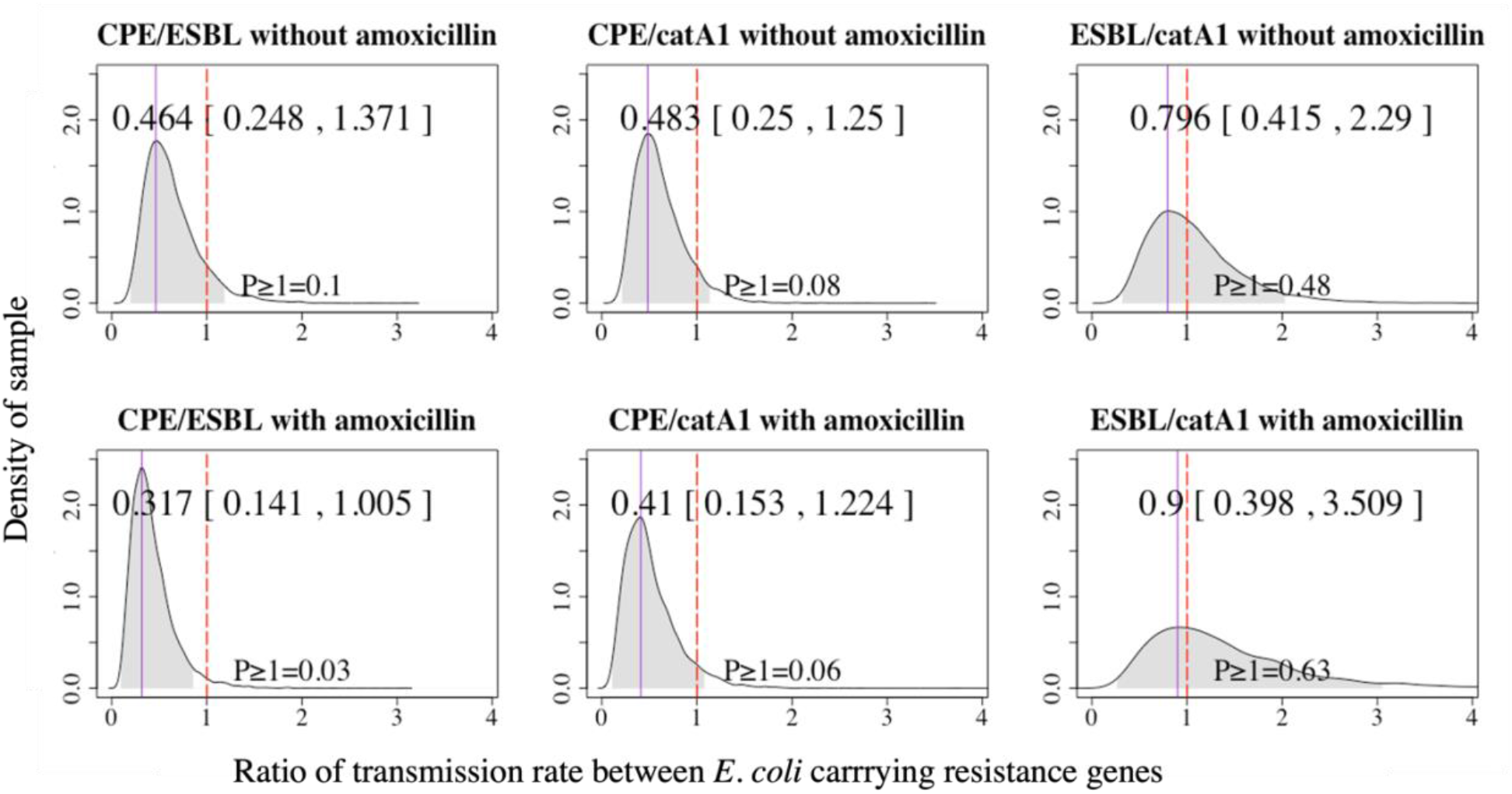
Density (vertical axis) of the posterior distribution of the ratio of the transmission rates (horizontal axis) for different inoculums: CPE-strain to ESBL-strain, CPE-strain to catA1-strain, and ESBL-strain to catA1-strain. The top and bottom row show plots for the pens without and with amoxicillin treatment, respectively. Purple vertical lines indicate the point estimate at the highest density and shaded areas are the 95% highest posterior density intervals of the posterior distribution; the estimated values of both are shown at the top of the plot. Dotted vertical red lines indicate a ratio of 1 and the probability of a ratio equal to or larger than 1 (P ≥ 1) is shown at the bottom of the plot.

The MAP of the estimated transmission rate of the CPE-strain is 46% and 48% of the transmission rate of the ESBL-strain and the catA1-strain in the non-amoxicillin-treated groups, respectively, and 32% and 41% of the transmission rate of ESBL-strain and catA1-strain in the amoxicillin-treated groups, respectively (Figure 5). HPDIs of the ratio of the transmission rates indicate the probability that transmission of the CPE-strain is faster than the transmission of the ESBL-strain or catA1-strain is 8% – 10% in non-amoxicillin-treated groups, and 3% – 6% in amoxicillin-treated groups (Figure 5). The MAP of the ratio of the ESBL-strain transmission rate to catA1-strain transmission rate is 0.80 without amoxicillin treatment and 0.90 with amoxicillin treatment, and the probability of a ratio equal to or larger than 1 is 0.48 and 0.63 for the groups without and with amoxicillin, respectively. This indicates the transmission rates of ESBL-strain and catA1-strain were similar in this experiment.

#### 3.2.3. Effect of amoxicillin

The transmission rates of all inoculums are smaller in the non-amoxicillin-treated groups than in the amoxicillin-treated groups (Figure 6). The difference between amoxicillin-treated groups and non-amoxicillin-treated groups is slightly larger for the ESBL-strain and catA1-strain than for the CPE-strain.

**Figure 6:**
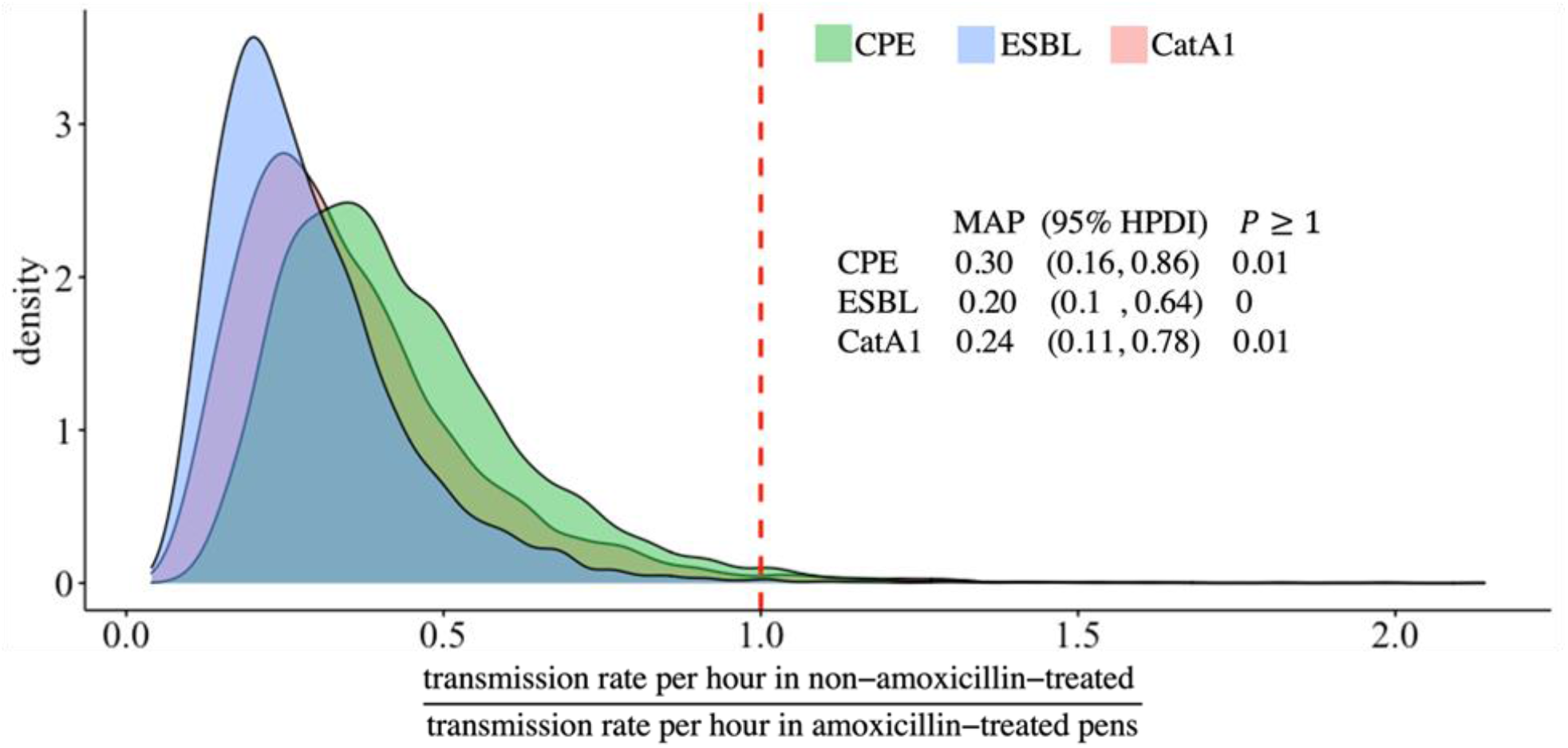
Density (vertical axis) of the ratio of the transmission rates in non-amoxicillin-treated pens over amoxicillin-treated pens (horizontal axis) for CPE-strain (green), ESBL-strain (blue) and catA1-strain (pink). The dotted red vertical line indicates a ratio of 1 (i.e., the transmission rates of amoxicillin-treated and non-amoxicillin groups are the same). The point estimate at the highest density (MAP) and 95% highest posterior density intervals (95% HPDI), and the probability of a ratio equal to or larger than 1 (P ≥ 1) are also shown in the plot.

### 3.3. Microbiome analysis

#### 3.3.1. Alpha-diversity

Observed richness measuring the observed number of taxa, Shannon’s index which takes evenness into account (with higher values if more taxa are present or taxa are more evenly distributed), and Pielou’s evenness which is not influenced by richness (with a value between 0 and 1, with higher values if taxa are more evenly distributed), were used to measure alpha-diversity. All alpha-diversity measures of the caecal microbiome at genus level on day 5 (i.e., before inoculation) were similar in the groups inoculated with the different inoculums (i.e., CPE-strain, ESBL-strain, catA1-strain; Figure 7). On day 14 various small differences in observed richness and Pielou’s evenness were found at genus level. Repeating these analyses at the level of individual ASVs mostly gave the same results (Figure S8; Tables S5 – S8).

**Figure 7.**
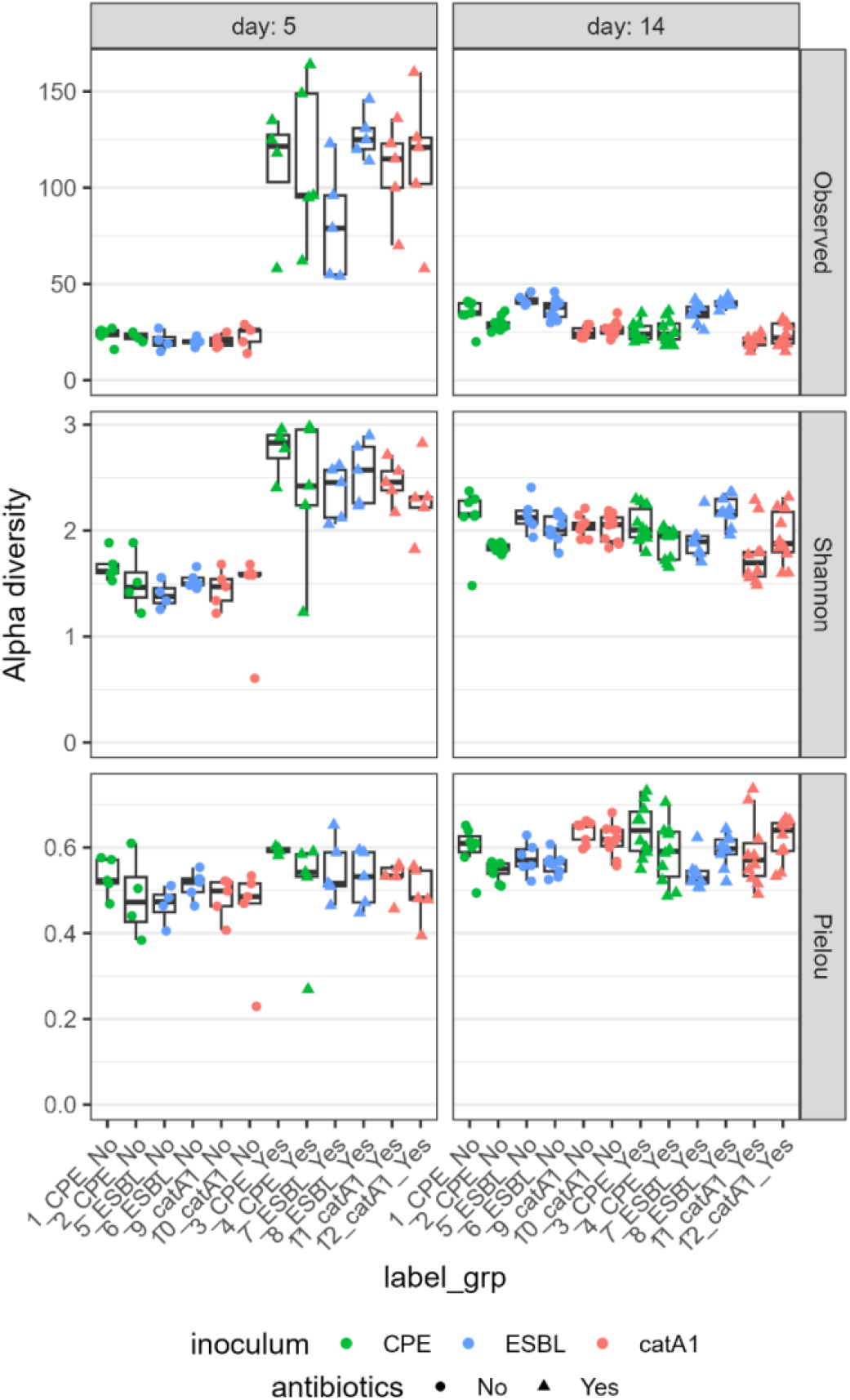
Alpha-diversity (vertical axis) by inoculum and antibiotic treatment (horizontal axis) at genus level. Colours indicate different inoculums (CPE-strain: green; ESBL-strain: blue; catA1-strain: red) and symbols indicate the absence (circles) or presence (triangles) of antibiotic treatment. The panels show the different alpha-diversity measures (rows) and different days (columns).

Observed richness and Shannon’s index at genus level on day 5 were lower in the non-amoxicillin-treated groups than in the amoxicillin-treated groups, but Pielou’s evenness was not different (Figure 7), indicating fewer genera were present in the non-amoxicillin-treated groups but the distribution of their abundances was similar to the distribution of their abundances in the amoxicillin-treated groups. By day 14, 8 days after finishing amoxicillin treatment, alpha-diversity was similar in the amoxicillin-treated and non-amoxicillin-treated groups. Repeating these analyses at the level of individual ASVs mostly gave the same results (Figure S8; Tables S5 – S8).

#### 3.3.2. Beta-diversity

The inoculums explained 6% and 3% of the variation between the groups in Bray-Curtis dissimilarity and Jaccard distance at genus level on day 5, antibiotic treatment explained 27% and 50% of the variation, and their interaction explained 5% and 3% of the variation (Table S9). Only groups without and with antibiotics were separated in the PCoA-plot (Figure 8). Repeating these analyses at the level of individual ASVs mostly gave the same results (see sections 2.5 and 2.6 of the supplementary material). Similarity percentage analyses showed the Bray-Curtis dissimilarities on day 5 between groups without and with antibiotic treatment are driven by the same genera in the groups inoculated with the different inoculation strains. Most of these genera belonged to the classes Bacilli and Clostridia, and some to the class Gammaproteobacteria (Tables S11 – S13).

**Figure 8.**
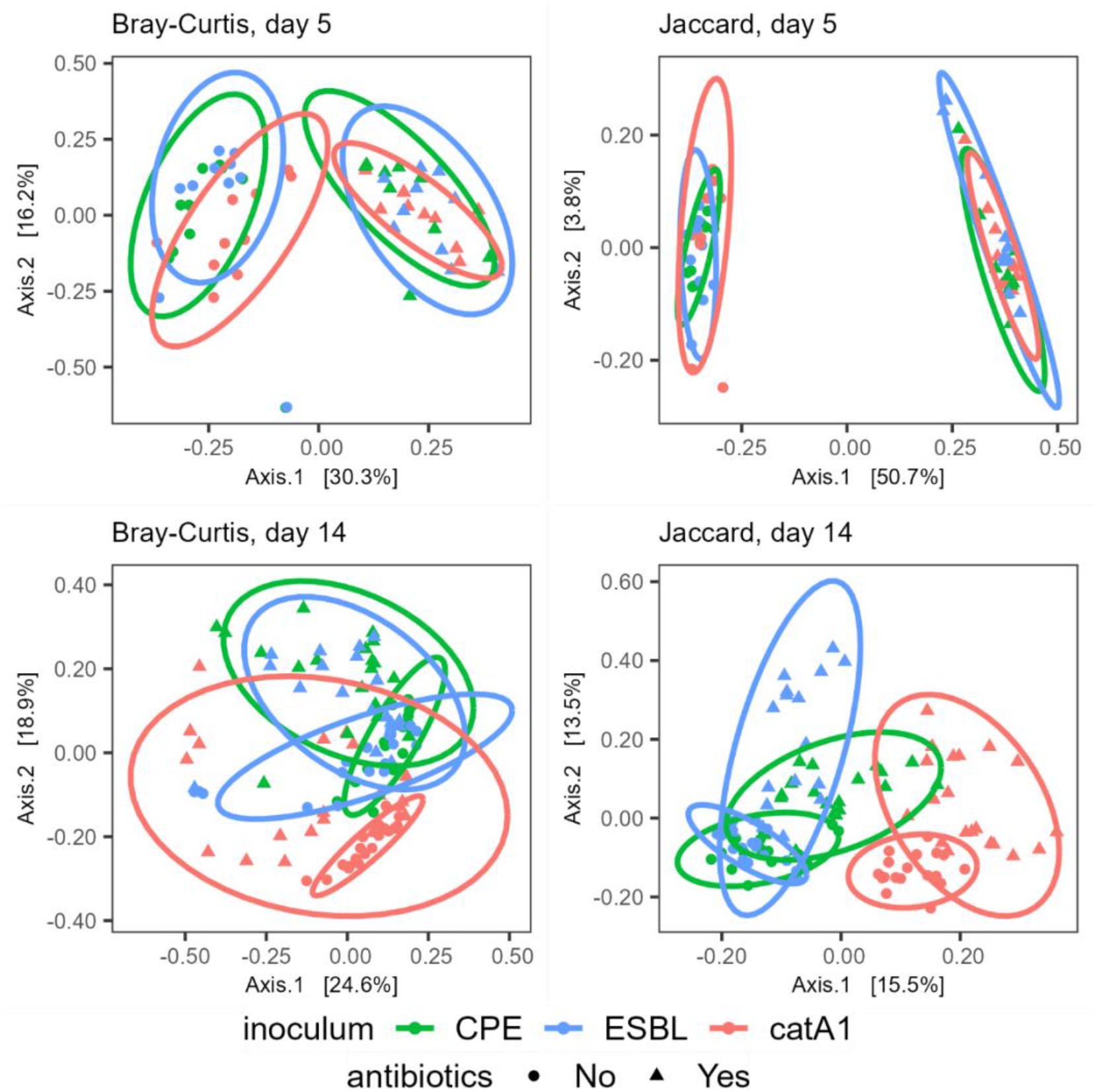
Principal coordinate plots based on Bray-Curtis dissimilarity (left) and Jaccard distance (right) for day 5 (top) and day 14 (bottom) at genus level. Colours indicate different inoculums (CPE-strain: green; ESBL-strain: blue; catA1-strain: red) and symbols indicate the absence (circles) or presence (triangles) of antibiotic treatment. Ellipses represent 95% confidence regions assuming a multivariate t-distribution.

The inoculums explained 16% and 17% of the variation between the groups in Bray-Curtis dissimilarity and Jaccard distance at genus level on day 14, antibiotic treatment explained 9% of the variation for both measures, and their interaction explained 4% and 6% of the variation (Table S10). For both beta-diversity measures, CPE-strain and ESBL-strain overlapped much with each other in the PCoA-plots, whereas catA1 without antibiotics separated from CPE-strain and ESBL-strain without antibiotics. Groups without and with antibiotics were not separate from each other on genus level (Figure 8) but separated on ASV level with Bray-Curtis dissimilarity (Figure S9).

## 4. Discussion

To our knowledge, this is the first transmission experiment with CPE *E. coli* in livestock. In addition, although the use of a Bayesian hierarchical model as presented in this study is well-recognized in epidemiology, its use in analysing animal transmission experiments is not common (Hu et al., 2017). Furthermore, we extended previous work on the relationship between the microbiome and the transmission of intestinal antibiotic-resistant bacteria (Dame-Korevaar et al., 2020b).

### 4.1. Indirect environmental transmission

*E. coli* is an enteric bacterium that is excreted in faeces and propagated in the environment (Conway and Cohen, 2015; Ramos et al., 2020), from where it can spread to other animals and humans (Rwego et al., 2008; Hussain et al., 2017; Rousham et al., 2018; Lepper et al., 2022). The environment can serve as a reservoir for the transmission of resistant bacteria when no excreting animals are present anymore (Dame-Korevaar et al., 2017). Therefore, we adapted the likelihood function to reflect environmental transmission with its prolonged possibility of transmission from accumulated bacteria in the environment.

It was not possible to estimate the decay rate and the transmission rate parameter with the Bayesian model simultaneously in our study, because a given number of cases can be explained equally well by a higher transmission rate or a lower decay rate. We reviewed the literature on decay rates (Table S14) to find a suitable range of decay rates and ran the hierarchical model with several fixed decay rates ranging from 0.04 – 55 h^-1^ (Table S15). This entire range of decay rates could be fitted well with low Watanabe–Akaike information criterion and divergence transition. Multiple studies in various environments suggest a very low level of *E. coli* decay in the first few days (see section 3.3 of the supplement), therefore we selected the lowest fixed decay rate (δ) of 0.04 h^-1^ in the final model.

The transmission rates of 3·10^−4^ h^-1^ and 1·10^−3^ h^-1^ for the ESBL-strain derived from our model assuming indirect environmental transmission are much lower than the transmission rate of 0.055 (0.045 – 0.066) h^-1^ calculated from a direct model (Dame-Korevaar et al., 2020b). A lower transmission rate is expected because resistant bacteria excreted into the environment were the only source of transmission considered in our SI-model and they decayed at a low rate. Using a higher decay rate would result in higher estimates for the transmission rates (Table S15), with a decay rate of 7.4 h^-1^ giving a transmission rate of 0.04 h^-1^ for ESBL without antibiotics, comparable to the value obtained by (Dame-Korevaar et al., 2020b). Using higher decay rates still results in transmission rates of CPE being lower than the transmission rates of the other inoculums.

### 4.2. Bayesian hierarchical inference

The Bayesian hierarchical model quantifies the transmission rate parameter of each pen using the mean transmission rate parameter and its variation simultaneously, instead of conventionally averaging the variation of all pens. This improves the estimates for each pen, especially when transmission events occur between sampling time points such that some pens have less information (McElreath, 2020). This was relevant for pens 7, 8, and 11 in which new cases were only observed at a very limited number of time points, because multiple transmission events occurred within the first few days (Figure 3), leading to wide HPDIs indicating a wide range of possible transmission rates. The hierarchical structure of the model led to shrinkage of the predicted data. Thus, we did not expect the predicted data to be equivalent to the observed data but instead expected systematic differences between the predicted and observed data (Figure 3).

The actual moment of transmission is rarely observed in transmission experiments because of logistic and ethical limitations to the number of animals and the sampling frequency (Cauchemez et al., 2004). A Bayesian approach in the analysis of transmission experiments can be used to incorporate the uncertainty that is inherent to the data in the statistical model, and to clearly present the uncertainty in the outcomes in the form of the posterior distribution (Hiura et al., 2021). These characteristics make Bayesian hierarchical modelling very suitable to quantify transmission between animals.

### 4.3. Effect of antibiotic resistance and *E. coli* strains on transmission

Resistance genes carried on plasmids generally impose fewer fitness costs on their bacterial hosts than chromosomal mutations resulting in resistance (Vogwill and MacLean, 2015). The fitness costs imposed by plasmids, lowering the population growth of resistant bacteria, can decrease when the number of plasmids within bacteria increases (Lee et al., 2020), increase when the number of resistance gene families on a plasmid increases and are also affected by host factors (Vogwill and MacLean, 2015). The CPE inoculum used in the animal experiment contained 3 plasmids carrying resistance genes from 6 families, while the ESBL strain contained 6 plasmids carrying resistance genes from 4 families, and the catA1 strain contained 4 plasmids carrying resistance genes from 4 families (Tables S16 – S18). The CPE-strain had more resistance gene families per plasmid and fewer plasmids than the other strains, both of which could increase fitness costs and thereby lower the transmission rate of CPE-strain compared to the other strains. On the other hand, in vitro conjugation experiments with multiple plasmids and resistance genes showed ESBL genes to be the costliest (Rajer and Sandegren, 2022) and the transmission rate of the CPE-strain was also lower in the presence of amoxicillin (Figure 5) when fitness costs are not expected to limit the transmission rate. This suggests the lower transmission rate of the CPE-strain is more likely caused by differences between the used *E. coli*-strains than by differences in plasmids and resistance genes.

The CPE, ESBL, and catA1 resistance genes used in the animal experiment were carried by different *E. coli* strains isolated from chickens between 2004 and 2009, so we cannot separate the effect of the different plasmids and the resistance genes they carried from the effect of the different *E. coli* strains. In addition, the resistance genes were located on conjugative plasmids and resistant colonies were not tested to identify the *E. coli* type. As such, part of the transmission might also be explained by plasmid transfer between *E. coli*, rather than by colonisation of the chicken gut by the *E. coli* strains that were present in the inoculums.

### 4.4. Effect of amoxicillin on transmission

Antibiotic treatment alleviates fitness costs of resistance genes and leads to resistant bacteria having a higher growth rate than susceptible bacteria, such that resistant bacteria would be expected to colonize the gut more easily and be transmitted faster in the amoxicillin-treated groups. Indeed, the transmission rates of all inoculums were higher in the amoxicillin-treated groups than in the non-amoxicillin-treated groups (Figure 6). Similarly, the relative abundance of the *E*.*coli/Shigella* genus was lower in amoxicillin-treated pens than in non-amoxicillin-treated pens on day 5 (i.e., before inoculation) but similar on day 14 (Figure S7), suggesting there was more ability for the inoculum to grow in antibiotic-treated pens. Nevertheless, the differences in transmission rates observed between the CPE-strain versus the ESBL-strain and the catA1-strain were also observed in amoxicillin-treated pens. This suggests intrinsic differences in the capability for transmission were present in these bacterial strains, which are independent of the antibiotic resistance itself, as we already stated above.

### 4.5. Microbiome analysis

Observed richness and evenness on day 14 differed between inoculums, but those differences were mainly caused by less within-group variation and were small compared to the effect of antibiotics on day 5 (Figure 7). The separation between the catA1-strain versus the CPE-strain and the ESBL-strain in beta-diversity on day 14 can be explained by broilers inoculated with the catA1-strain being housed in a room separate from broilers inoculated with the CPE-strain and ESBL-strain, in addition to the effect of being inoculated with a different *E. coli* strain. This room effect was also reflected in the caecal composition of the non-amoxicillin-treated catA1 groups being more similar to the composition of the amoxicillin-treated catA1 groups than to the composition of the non-amoxicillin-treated CPE-groups and ESBL-groups at family level (Figure S5).

The differences in alpha-diversity and beta-diversity between the different inoculums do not correspond to the differences in the transmission rates between the inoculums. The ESBL-strain contained slightly more observed taxa on day 14, the catA1-strain contained a slightly more even distribution of taxa on day 14, and the catA1-strain separated somewhat from CPE and ESBL on genus level and clearly on ASV level. The transmission rates of the ESBL-strain and the catA1-strain were, however, very similar and the transmission rate of the CPE-strain was lower. This indicates the differences in transmission between the inoculums are most likely not caused by differences in the caecal microbiome.

Observed richness on day 5 was lower in non-amoxicillin-treated pens than in amoxicillin-treated pens, but Pielou’s evenness was similar (Figure 7), suggesting amoxicillin treatment allowed more taxa to increase in abundance but did not lead to differences in the proportion of dominating taxa. The lower richness in non-amoxicillin-treated pens was the opposite of the higher richness expected based on the literature mentioned in the introduction, which might have been caused by the depletion of some major abundant taxa by the amoxicillin treatment, leaving more room for rare taxa to be detected by the sequencing depth that became available. Amoxicillin treatment did affect the microbiome composition at class, family level and genus level (Figure S4; Figure S5; Figure S6). On day 14, only few differences in alpha-diversity and evenness between the non-amoxicillin-treated and amoxicillin-treated groups were found (Figure 7). Similarity percentage analyses indicated the effects of antibiotic treatment on Bray-Curtis dissimilarity on day 5 were driven by the same genera in the groups inoculated with the different inoculation strains (Tables S11 – S13). Amoxicillin treatment explained less variation in beta-diversity on day 14 than on day 5, and the non-amoxicillin-treated and amoxicillin-treated groups did not separate clearly in the PCoA plot at genus level on day 14. This indicates differences in the genera present in the caecal microbiome on day 5 caused by antibiotic treatment did not last until day 14. Amoxicillin is cleared quickly from chickens when administration ceases and decays quickly in the environment (Peng et al., 2016), such that the effect of amoxicillin might have been reduced by day 14 because it was last administered on day 6. Although other clinically important antibiotics such as cephalosporins are cleared slower and could last longer in the environment such that they could have an effect on day 14, we did not incorporate them in our study because their use in livestock is subject to legal restrictions (Bonten et al., 2021). The higher alpha-diversity in amoxicillin-treated groups on day 5 might be related to the higher transmission rates in amoxicillin-treated groups because transmission events mainly occurred within a few days after inoculation (Figure 3). The microbiome of broilers evolves in steps to a more or less stable state in 35 days (Jurburg et al., 2019; Kers et al., 2022). We hypothesize that the dysbiosis of the microbiome caused by antibiotic treatment allows for easier colonization and more rapid growth of new *E. coli* strains such as the inoculums, which is reflected in a more rapid transmission. The opposite, e.g., quicker maturation of the gut microbiome by applying a probiotic, has been shown to slow down transmission (Ceccarelli et al., 2017; Dame-Korevaar et al., 2020b).

### 4.6. Suggestions for further research

The uncertainty and variability of the transmission rates of the 3 *E. coli* strains provide a good range of transmission rates needed to model the transmission of resistance genes carried by commensal bacteria in poultry. Future research could expand the Bayesian hierarchical framework adopted in this study by incorporating data together with their uncertainty from other experiments on bacterial transmission between broilers to capture the influence of differences in environments, chickens’ feed, and different species of resistant bacteria. This would result in a transmission model that reflects the situation on broiler farms more realistically.

In a clean environment, inoculated broilers should start excreting before contact broilers can be colonized. However, in some pens in this experiment, the first excretion of resistant bacteria by both inoculated and contact broilers was detected at the same sampling time point. This is caused by limitations to the sampling frequency. We could use the model by assuming that inoculated broilers started excreting half a time interval earlier. This assumption has previously been used in the analysis of a transmission experiment in broilers where the moment of excretion was similar for inoculated and contact animals (Dame-Korevaar et al., 2020a). In future research, estimation of the exact time point of colonization could be incorporated, e.g., by applying the Bayesian approach described for a model of direct transmission (Hu et al., 2017) to a model of environmental transmission. Taking more frequent samples could also help, although more frequent sampling of the caeca is limited by ethical considerations.

Although the presence of multiple plasmids in a bacterium reflects the situation that is common in nature (Davies and Davies, 2010; MacLean and San Millan, 2015), future research should compare the transmission rates of different resistance genes using a single *E. coli* strain that only contains the plasmid of interest for the different inoculums. We were not able to use that approach because of a lack of the necessary permits to work with genetically modified organisms in animal experiments, but here we showed the difference in transmission rates between strains could be substantial (up to 68%) and is thus relevant. Using that same *E. coli* strain with chromosomal resistance instead of plasmids as inoculum would allow for the comparison of the transmission of plasmid-mediated and chromosomal resistance. Such research can build on this paper by applying the same methodology and determining their sampling schemes based on our results.

## 5. Summary

From our study, we conclude early amoxicillin treatment increases the transmission rate of *E. coli* strains carrying different resistance genes between broilers up to five-fold. Amoxicillin treatment increased alpha-diversity of the caecal microbiome on day 5, but no effects of amoxicillin treatment on the caecal microbiome were found on day 14, suggesting it only has a temporary effect on the caecal microbiome. In addition, the effects of amoxicillin on the transmission rates were most likely not caused by differences in the caecal microbiome, because differences in the microbiomes of the different inoculums did not correspond to the differences in the transmission rates of the different inoculums. The transmission rates of 2·10^−4^ h^-1^ and 4·10^−4^ h^-1^ for the CPE-strain were 54 – 68 per cent lower than the transmission rates of the ESBL-strain and 52 – 59 per cent lower than the transmission rates of the catA1-strain. This was reflected in the longer time needed for the CPE-strain to colonize all broilers than for the ESBL-strain and catA1-strain. Such delays might be relevant in the field, especially if competition between different antibiotic-resistant strains occurs. The consistent difference in transmission rates with and without antibiotic treatment indicates the differences in transmission rates were more likely caused by differences between the used *E. coli* strains than by differences in plasmids and resistance genes. The methodology applied in this experiment and the obtained transmission rates can be used to improve models of the transmission of resistant bacteria between broilers.

## Abbreviations

ASV: amplicon sequence variant
BSL: Biosafety level
CPE: carbapenemase-producing *Enterobacteriaceae*
*E. coli*: *Escherichia coli*
ESBL: extended-spectrum beta-lactamase
HPDI: highest posterior density interval
MAP: maximum a posteriori estimate
PCoA: Principal coordinate analysis
SI-model: susceptible-infectious model
WBVR: Wageningen Bioveterinary Research

## 6. Acknowledgements

The authors gratefully acknowledge the assistance provided during this project by the animal caretakers, pathologists and technicians at Wageningen Bioveterinary Research.

## 7. Funding

This work was supported by ZonMW [grant numbers 50-54100-98-229, 50-54100-98-119].

## Notes

### Competing Interest Statement

The authors have declared no competing interest.

## References

Aarestrup, F.M., 2015. The livestock reservoir for antimicrobial resistance: a personal view on changing patterns of risks, effects of interventions and the way forward. Philos Trans R Soc Lond B Biol Sci 370, 20140085.

Bonardi, S., Pitino, R., 2019. Carbapenemase-producing bacteria in food-producing animals, wildlife and environment: a challenge for human health. Ital J Food Saf 8, 7956–7956.

Bonten, M.J.M., van Geijlswijk, I.M., Heederik, D.J.J., Mevius, D.J. P. S., 2021. Usage of antibiotics in agricultural livestock in the Netherlands in 2020 - appendix.

Bortolaia, V., Ronco, T., Romascu, L., Nicorescu, I., Milita, N.M., Vaduva, A.M., Leekitcharoenphon, P., Kjeldgaard, J.S., Hansen, I.M., Svendsen, C.A., Mordhorst, H., Guerra, B., Beloeil, P.A., Hoffmann, M., Hendriksen, R.S., 2021. Co-localization of carbapenem (blaOXA-162) and colistin (mcr-1) resistance genes on a transferable IncHI2 plasmid in Escherichia coli of chicken origin. J Antimicrob Chemother 76, 3063–3065.

Brink, A.J., 2019. Epidemiology of carbapenem-resistant Gram-negative infections globally. Curr Opin Infect Dis 32, 609–616.

Callahan, B.J., McMurdie, P.J., Rosen, M.J., Han, A.W., Johnson, A.J.A., Holmes, S.P., 2016. DADA2: High-resolution sample inference from Illumina amplicon data. Nature Methods 13, 581–583.

Cauchemez, S., Carrat, F., Viboud, C., Valleron, A.J., Boëlle, P.Y., 2004. A Bayesian MCMC approach to study transmission of influenza: application to household longitudinal data. Stat Med 23, 3469–3487.

Ceccarelli, D., van Essen-Zandbergen, A., Smid, B., Veldman, K.T., Boender, G.J., Fischer, E.A.J., Mevius, D.J., van der Goot, J.A., 2017. Competitive exclusion reduces transmission and excretion of extended-spectrum-β-lactamase-producing Escherichia coli in broilers. Appl Environ Microbiol 83.

Conway, T., Cohen, P.S., 2015. Commensal and pathogenic Escherichia coli metabolism in the gut. Microbiol Spectr 3.

Dame-Korevaar, A., Fischer, E.A.J., Stegeman, A., Mevius, D., van Essen-Zandbergen, A., Velkers, F., van der Goot, J., 2017. Dynamics of CMY-2 producing E. coli in a broiler parent flock. Vet Microbiol 203, 211–214.

Dame-Korevaar, A., Fischer, E.A.J., van der Goot, J., Velkers, F., Ceccarelli, D., Mevius, D., Stegeman, A., 2020a. Early life supply of competitive exclusion products reduces colonization of extended spectrum beta-lactamase-producing Escherichia coli in broilers. Poult Sci 99, 4052–4064.

Dame-Korevaar, A., Fischer, E.A.J., van der Goot, J., Velkers, F., van den Broek, J., Veldman, K., Ceccarelli, D., Mevius, D., Stegeman, A., 2019. Effect of challenge dose of plasmid-mediated extended-spectrum β-lactamase and AmpC β-lactamase producing Escherichia coli on time-until-colonization and level of excretion in young broilers. Vet Microbiol 239, 108446.

Dame-Korevaar, A., Kers, J.G., van der Goot, J., Velkers, F.C., Ceccarelli, D., Mevius, D.J., Stegeman, A., Fischer, E.A.J., 2020b. Competitive exclusion prevents colonization and compartmentalization reduces transmission of ESBL-producing Escherichia coli in broilers. Front Microbiol 11, 566619.

Dankittipong, N., Fischer, E.A.J., Swanenburg, M., Wagenaar, J.A., Stegeman, A.J., de Vos, C.J., 2022. Quantitative risk assessment for the introduction of carbapenem-resistant Enterobacteriaceae (CPE) into dutch livestock farms. Antibiotics (Basel) 11.

Davies, J., Davies, D., 2010. Origins and evolution of antibiotic resistance. Microbiol Mol Biol Rev 74, 417–433.

Dinno, A., 2017. dunn.test: Dunn’s test of multiple comparisons using rank sums.

European Centre for Disease Prevention and Control, 2018. Surveillance of antimicrobial resistance in Europe – Annual report of the European Antimicrobial Resistance Surveillance Network (EARS-Net) 2017. Stockholm.

European Food Safety Authority, European Centre for Disease Prevention Control, 2022. The European Union summary report on antimicrobial resistance in zoonotic and indicator bacteria from humans, animals and food in 2019–2020. EFSA Journal 20, e07209.

Finotello, F., Mastrorilli, E., Di Camillo, B., 2018. Measuring the diversity of the human microbiota with targeted next-generation sequencing. Brief Bioinform 19, 679–692.

Gabry, J., Cešnovar, R., 2022. cmdstanr: R Interface to ‘CmdStan’.

Gerhards, N.M., Gonzales, J.L., Vreman, S., Ravesloot, L., van den Brand, J.M.A., Doekes, H.P., Egberink, H.F., Stegeman, A., Oreshkova, N., van der Poel, W.H.M., de Jong, M.C.M., 2022. Efficient direct and limited environmental transmission of SARS-CoV-2 lineage B.1.22 in domestic cats. bioRxiv, 2022.2006.2017.496600.

Hiura, S., Abe, H., Koyama, K., Koseki, S., 2021. Bayesian generalized linear model for simulating bacterial inactivation/growth considering variability and uncertainty. Front Microbiol 12, 674364.

Holmes, A.H., Moore, L.S.P., Sundsfjord, A., Steinbakk, M., Regmi, S., Karkey, A., Guerin, P.J., Piddock, L.J.V., 2016. Understanding the mechanisms and drivers of antimicrobial resistance. The Lancet 387, 176–187.

Hu, B., Gonzales, J.L., Gubbins, S., 2017. Bayesian inference of epidemiological parameters from transmission experiments. Sci Rep 7, 16774.

Huijbers, P.M.C., Graat, E.A.M., van Hoek, A., Veenman, C., de Jong, M.C.M., van Duijkeren, E., 2016. Transmission dynamics of extended-spectrum β-lactamase and AmpC β-lactamase-producing Escherichia coli in a broiler flock without antibiotic use. Prev Vet Med 131, 12–19.

Hussain, A., Shaik, S., Ranjan, A., Nandanwar, N., Tiwari, S.K., Majid, M., Baddam, R., Qureshi, I.A., Semmler, T., Wieler, L.H., Islam, M.A., Chakravortty, D., Ahmed, N., 2017. Risk of transmission of antimicrobial resistant Escherichia coli from commercial broiler and free-range retail chicken in India. Front Microbiol 8, 2120.

Jurburg, S.D., Brouwer, M.S.M., Ceccarelli, D., van der Goot, J., Jansman, A.J.M., Bossers, A., 2019. Patterns of community assembly in the developing chicken microbiome reveal rapid primary succession. Microbiologyopen 8, e00821.

Keeling, M.J., Rohani, P., 2007. Modeling Infectious Diseases in Humans and Animals Princeton University Press.

Kers, J.G., Velkers, F.C., Fischer, E.A.J., Stegeman, J.A., Smidt, H., Hermes, G.D.A., 2022. Conserved developmental trajectories of the cecal microbiota of broiler chickens in a field study. FEMS Microbiology Ecology 98.

Kim, S., Covington, A., Pamer, E.G., 2017. The intestinal microbiota: antibiotics, colonization resistance, and enteric pathogens. Immunol Rev 279, 90–105.

Knobler, S., Marian, N., Stacey, L.K., Stanley, M.L., Tom, B., Health, B.o.G., Burroughs, T., Infections, F.o.E., Knobler, S.L., Lemon, S.M., Najafi, M., 2003. The resistance phenomenon in microbes and infectious disease vectors: implications for human health and strategies for containment (workshop summary). National Academies Press 2003.

Köck, R., Daniels-Haardt, I., Becker, K., Mellmann, A., Friedrich, A.W., Mevius, D., Schwarz, S., Jurke, A., 2018. Carbapenem-resistant Enterobacteriaceae in wildlife, food-producing, and companion animals: a systematic review. Clin Microbiol Infect 24, 1241–1250.

Lahti, L., Shetty, S., 2019. microbiome R package.

Lee, H., Shin, J., Chung, Y.J., Park, M., Kang, K.J., Baek, J.Y., Shin, D., Chung, D.R., Peck, K.R., Song, J.H., Ko, K.S., 2020. Co-introduction of plasmids harbouring the carbapenemase genes, bla(NDM-1) and bla(OXA-232), increases fitness and virulence of bacterial host. J Biomed Sci 27, 8.

Leekitcharoenphon, P., Johansson, M.H.K., Munk, P., Malorny, B., Skarżyńska, M., Wadepohl, K., Moyano, G., Hesp, A., Veldman, K.T., Bossers, A., Zając, M., Wasyl, D., Sanders, P., Gonzalez-Zorn, B., Brouwer, M.S.M., Wagenaar, J.A., Heederik, D.J.J., Mevius, D., Aarestrup, F.M., 2021. Genomic evolution of antimicrobial resistance in Escherichia coli. Sci Rep 11, 15108.

Lepper, H.C., Woolhouse, M.E.J., van Bunnik, B.A.D., 2022. The role of the environment in dynamics of antibiotic resistance in humans and animals: a modelling study. Antibiotics.

Leverstein-van Hall, M.A., Dierikx, C.M., Cohen Stuart, J., Voets, G.M., van den Munckhof, M.P., van Essen-Zandbergen, A., Platteel, T., Fluit, A.C., van de Sande-Bruinsma, N., Scharinga, J., Bonten, M.J., Mevius, D.J., 2011. Dutch patients, retail chicken meat and poultry share the same ESBL genes, plasmids and strains. Clin Microbiol Infect 17, 873–880.

Lister, S.A., Barrow, P., 2008. Chapter 8 - Enterobacteriaceae. In: Pattison, M., McMullin, P.F., Bradbury, J.M., Alexander, D.J. (Eds.), Poultry diseases (sixth edition). W.B. Saunders, Edinburgh, 110–145.

MacLean, R.C., San Millan, A., 2015. Microbial evolution: towards resolving the plasmid paradox. Curr Biol 25, R764–767.

Madec, J.Y., Haenni, M., 2018. Antimicrobial resistance plasmid reservoir in food and food-producing animals. Plasmid 99, 72–81.

Makowski, D., Ben-Shachar, M.S., Lüdecke, D., 2019. bayestestR: describing effects and their uncertainty, existence and significance within the Bayesian framework. Journal of open source software 4.

Mathew, A.G., Cissell, R., Liamthong, S., 2007. Antibiotic resistance in bacteria associated with food animals: a United States perspective of livestock production. Foodborne Pathog Dis 4, 115–133.

McElreath, R., 2020. Statistical rethinking: a bayesian course with examples in R and Stan. Chapman & Hall.

McMurdie, P.J., Holmes, S., 2013. phyloseq: An R package for reproducible interactive analysis and graphics of microbiome census data. PLOS ONE 8, e61217.

Mughini-Gras, L., Dorado-García, A., van Duijkeren, E., van den Bunt, G., Dierikx, C.M., Bonten, M.J.M., Bootsma, M.C.J., Schmitt, H., Hald, T., Evers, E.G., de Koeijer, A., van Pelt, W., Franz, E., Mevius, D.J., Heederik, D.J.J., 2019. Attributable sources of community-acquired carriage of Escherichia coli containing β-lactam antibiotic resistance genes: a population-based modelling study. Lancet Planet Health 3, e357–e369.

Nielsen, L.R., van den Borne, B., van Schaik, G., 2007. Salmonella Dublin infection in young dairy calves: transmission parameters estimated from field data and an SIR-model. Prev Vet Med 79, 46–58.

Ogunrinu, O.J., Norman, K.N., Vinasco, J., Levent, G., Lawhon, S.D., Fajt, V.R., Volkova, V.V., Gaire, T., Poole, T.L., Genovese, K.J., Wittum, T.E., Scott, H.M., 2020. Can the use of older-generation beta-lactam antibiotics in livestock production over-select for beta-lactamases of greatest consequence for human medicine? An in vitro experimental model. PLoS One 15, e0242195.

Oksanen, J., Simpson, G.L., Blanchet, F.G., Kindt, R., Legendre, P., Minchin, P.R., O’Hara, R.B., Solymos, P., Stevens, M.H.H., Szoecs, E., Wagner, H., Barbour, M., Bedward, M., Bolker, B., Borcard, D., Carvalho, G., Chirico, M., De Caceres, M., Durand, S., Evangelista, H.B.A., FitzJohn, R., Friendly, M., Furneaux, B., Hannigan, G., Hill, M.O., Leo Lahti, L., McGlinn, D., Ouellette, M., Cunha, E.R., Smith, T., Stier, A., Ter Braak, C.J.F., Weedon, J., 2022. Vegan: community ecology package.

Pagès, H., Aboyoun P., Gentleman R., DebRoy S., 2022. Biostrings: efficient manipulation of biological strings.

Peng, P.C., Wang, Y., Liu, L.Y., Zou, Y.D., Liao, X.D., Liang, J.B., Wu, Y.B., 2016. The excretion and environmental effects of amoxicillin, ciprofloxacin, and doxycycline residues in layer chicken manure. Poult Sci 95, 1033–1041.

Quast, C., Pruesse, E., Yilmaz, P., Gerken, J., Schweer, T., Yarza, P., Peplies, J., Glöckner, F.O., 2012. The SILVA ribosomal RNA gene database project: improved data processing and web-based tools. Nucleic Acids Research 41, D590–D596.

R Core Team, 2020. R: A language and environment for statistical computing. R Foundation for Statistical Computing, Vienna, Austria.

R Core Team, 2021. R: A language and environment for statistical computing. R Foundation for Statistical Computing, Vienna, Austria.

Rajer, F., Sandegren, L., 2022. The role of antibiotic resistance genes in the fitness cost of multiresistance plasmids. mBio 13, e0355221.

Ramos, S., Silva, V., Dapkevicius, M.L.E., Caniça, M., Tejedor-Junco, M.T., Igrejas, G., Poeta, P., 2020. Escherichia coli as commensal and pathogenic bacteria among food-producing animals: health implications of extended spectrum β-lactamase (ESBL) production. Animals (Basel) 10.

Robé, C., Blasse, A., Merle, R., Friese, A., Roesler, U., Guenther, S., 2019. Low dose colonization of broiler chickens with ESBL-/AmpC-producing Escherichia coli in a seeder-bird model independent of antimicrobial selection pressure. Front Microbiol 10, 2124.

Rochegüe, T., Haenni, M., Mondot, S., Astruc, C., Cazeau, G., Ferry, T., Madec, J.-Y., Lupo, A., 2021. Impact of antibiotic therapies on resistance genes dynamic and composition of the animal gut microbiota. Animals 11, 3280.

Rousham, E.K., Unicomb, L., Islam, M.A., 2018. Human, animal and environmental contributors to antibiotic resistance in low-resource settings: integrating behavioural, epidemiological and One Health approaches. Proceedings of the Royal Society B: Biological Sciences 285, 20180332.

Rozwandowicz, M., Brouwer, M.S.M., Fischer, J., Wagenaar, J.A., Gonzalez-Zorn, B., Guerra, B., Mevius, D.J., Hordijk, J., 2018. Plasmids carrying antimicrobial resistance genes in Enterobacteriaceae. J Antimicrob Chemother 73, 1121–1137.

Rwego, I.B., Gillespie, T.R., Isabirye-Basuta, G., Goldberg, T.L., 2008. High rates of Escherichia coli transmission between livestock and humans in rural Uganda. J Clin Microbiol 46, 3187–3191.

Schmidt, T.S., Matias Rodrigues, J.F., von Mering, C., 2017. A family of interaction-adjusted indices of community similarity. Isme j 11, 791–807.

Sorbara, M.T., Pamer, E.G., 2019. Interbacterial mechanisms of colonization resistance and the strategies pathogens use to overcome them. Mucosal Immunol 12, 1–9.

Stan Development Team, 2018. StanHeaders: headers for the R interface to Stan.

Stan Development Team, 2020. RStan: the R interface to Stan.

van Bunnik, B.A., Ssematimba, A., Hagenaars, T.J., Nodelijk, G., Haverkate, M.R., Bonten, M.J., Hayden, M.K., Weinstein, R.A., Bootsma, M.C., De Jong, M.C., 2014. Small distances can keep bacteria at bay for days. Proc Natl Acad Sci U S A 111, 3556–3560.

van Elsas, J.D., Semenov, A.V., Costa, R., Trevors, J.T., 2011. Survival of Escherichia coli in the environment: fundamental and public health aspects. Isme j 5, 173–183.

Velthuis, A.G., Bouma, A., Katsma, W.E., Nodelijk, G., De Jong, M.C., 2007. Design and analysis of small-scale transmission experiments with animals. Epidemiol Infect 135, 202–217.

Ventola, C.L., 2015. The antibiotic resistance crisis: part 1: causes and threats. P t 40, 277–283.

Vogwill, T., MacLean, R.C., 2015. The genetic basis of the fitness costs of antimicrobial resistance: a meta-analysis approach. Evol Appl 8, 284–295.

Wickham, H., 2016. ggplot2: elegant graphics for data analysis. Springer-Verlag, New York.

Wickham, H., François, R., Henry, L., Müller, K., 2021. dplyr: a grammar of data manipulation.

Wickham, H., Girlich, M., 2022. tidyr: tidy messy data.

Wilke, C.O., 2020. cowplot: streamlined plot theme and plot annotations for ‘ggplot2’.

Wilson, H., Török, M.E., 2018. Extended-spectrum β-lactamase-producing and carbapenemase-producing Enterobacteriaceae. Microb Genom 4.

World Health Organization, 2019. Critically important antimicrobials for human medicine, 6th revision. World Health Organization, Geneva.

Wu, G., Day, M.J., Mafura, M.T., Nunez-Garcia, J., Fenner, J.J., Sharma, M., van Essen-Zandbergen, A., Rodríguez, I., Dierikx, C., Kadlec, K., Schink, A.K., Chattaway, M., Wain, J., Helmuth, R., Guerra, B., Schwarz, S., Threlfall, J., Woodward, M.J., Woodford, N., Coldham, N., Mevius, D., 2013. Comparative analysis of ESBL-positive Escherichia coli isolates from animals and humans from the UK, The Netherlands and Germany. PLoS One 8, e75392.

Zhou, R., Fang, X., Zhang, J., Zheng, X., Shangguan, S., Chen, S., Shen, Y., Liu, Z., Li, J., Zhang, R., Shen, J., Walsh, T.R., Wang, Y., 2021. Impact of carbapenem resistance on mortality in patients infected with Enterobacteriaceae: a systematic review and meta-analysis. BMJ Open 11, e054971.

