## Supplementary material for "Comparing the transmission of carbapenemase-producing and extended-spectrum beta-lactamase-producing *Escherichia coli* between broiler chickens"

### 1. Data from the transmission experiment

#### 1.1. Raw transmission data

The pen, used inoculum, antibiotic treatment, and the test results of the cloacal swabs (i.e., positive or negative for CPE-strain, ESBL-strain, or catA1-strain) at each sampling time point were recorded for all inoculated and contact broilers (Table S1).

*Table S1: The experimental treatments and status at each sampling time point for all inoculated and contact broilers. Light shading indicates a positive test result, dark shading indicates the absence of a test result because the broiler died. Abbreviations: AMU: antimicrobial usage.*

| | | | | Sampling time point (hours after inoculation) | | | | | | | | | |
|---|---|---|---|---|---|---|---|---|---|---|---|---|---|
| Pen | Inoculum | AMU | ID | 0 | 8 | 24 | 32 | 48 | 72 | 96 | 120 | 168 | 216 |
| 1 | CPE | - | 0101 | 0 | 0 | 1 | D | D | D | D | D | D | D |
| 1 | CPE | - | 0114 | 0 | 0 | 1 | 0 | 1 | 1 | 1 | 1 | 1 | 1 |
| 1 | CPE | - | 0110 | 0 | 0 | 0 | 0 | 1 | 1 | 1 | 1 | 1 | 1 |
| 1 | CPE | - | 0108 | 0 | 0 | 0 | 0 | 0 | 1 | 0 | 1 | 1 | 1 |
| 1 | CPE | - | 0109 | 0 | 0 | 0 | D | D | D | D | D | D | D |
| 1 | | - | 0113 | 0 | 0 | 0 | 0 | 0 | 1 | 0 | 1 | 1 | 1 |
| 1 | | - | 0102 | 0 | 0 | 0 | 0 | 0 | 0 | 1 | 1 | 1 | 1 |
| 1 | | - | 0103 | 0 | 0 | 0 | 0 | 0 | 0 | 1 | 0 | 1 | 1 |
| 1 | | - | 0104 | 0 | 0 | 0 | 0 | 0 | 0 | 1 | 0 | 1 | 1 |
| 1 | | - | 0107 | 0 | 0 | 0 | 0 | 0 | D | D | D | D | D |
| 2 | CPE | - | 0209 | 0 | 1 | 0 | 0 | 0 | 1 | 1 | 1 | 1 | 1 |
| 2 | CPE | - | 0202 | 0 | 0 | 0 | 1 | 1 | 1 | 1 | 1 | 1 | 1 |
| 2 | CPE | - | 0205 | 0 | 0 | 0 | 0 | 1 | 1 | 0 | 1 | 1 | 1 |
| 2 | CPE | - | 0211 | 0 | 0 | 0 | 0 | 0 | 1 | 1 | 0 | 1 | 1 |
| 2 | CPE | - | 0212 | 0 | 0 | 0 | D | D | D | D | D | D | D |
| 2 | | - | 0206 | 0 | 0 | 0 | 0 | 1 | 1 | 0 | 0 | 1 | 1 |
| 2 | | - | 0204 | 0 | 0 | 0 | 0 | 0 | 1 | 0 | 1 | 1 | 1 |
| 2 | | - | 0208 | 0 | 0 | 0 | 0 | 0 | 1 | 0 | 0 | 1 | 1 |
| 2 | | - | 0215 | 0 | 0 | 0 | 0 | 0 | 1 | 1 | 1 | 1 | 1 |
| 2 | | - | 0201 | 0 | 0 | 0 | 0 | 0 | 0 | 0 | 0 | 1 | 1 |
| 3 | CPE | + | 0304 | 0 | 0 | 0 | 0 | 0 | 1 | 1 | 1 | 1 | 1 |
| 3 | CPE | + | 0310 | 0 | 0 | 0 | 0 | 0 | 0 | 1 | 1 | 1 | 1 |

| | | | | | | | | | | | | | |
|---|---|---|---|---|---|---|---|---|---|---|---|---|---|
| 3 | CPE | + | 0312 | 0 | 0 | 0 | 0 | 0 | 0 | 1 | 1 | 1 | 1 |
| 3 | CPE | + | 0303 | 0 | 0 | 0 | 0 | 0 | 0 | 0 | 1 | 1 | 1 |
| 3 | CPE | + | 0313 | 0 | 0 | 0 | 0 | 0 | 0 | 0 | 0 | 1 | 1 |
| 3 | | + | 0308 | 0 | 0 | 0 | 0 | 0 | 1 | 0 | 1 | 1 | 1 |
| 3 | | + | 0306 | 0 | 0 | 0 | 0 | 0 | 0 | 1 | 1 | 1 | 1 |
| 3 | | + | 0307 | 0 | 0 | 0 | 0 | 0 | 0 | 1 | 1 | 1 | 1 |
| 3 | | + | 0315 | 0 | 0 | 0 | 0 | 0 | 0 | 0 | 1 | 1 | 1 |
| 3 | | + | 0309 | 0 | 0 | 0 | 0 | 0 | 0 | 0 | 0 | 1 | 1 |
| 4 | CPE | + | 0402 | 0 | 0 | 1 | 1 | 1 | 1 | 1 | 1 | 1 | 1 |
| 4 | CPE | + | 0404 | 0 | 0 | 1 | 0 | 1 | 1 | 1 | 1 | 1 | 1 |
| 4 | CPE | + | 0405 | 0 | 0 | 1 | 1 | 1 | 1 | 1 | 1 | 1 | 1 |
| 4 | CPE | + | 0406 | 0 | 0 | 1 | 1 | 1 | 1 | 1 | 1 | 1 | 1 |
| 4 | CPE | + | 0415 | 0 | 0 | 1 | 1 | 1 | 1 | 1 | 1 | 1 | 1 |
| 4 | | + | 0401 | 0 | 0 | 1 | 1 | 1 | 1 | 1 | 1 | 1 | 1 |
| 4 | | + | 0409 | 0 | 0 | 1 | 1 | 1 | 1 | 1 | 1 | 1 | 1 |
| 4 | | + | 0413 | 0 | 0 | 1 | 1 | 1 | 1 | 1 | 1 | 1 | 1 |
| 4 | | + | 0407 | 0 | 0 | 0 | 1 | 1 | 1 | 1 | 1 | 1 | 1 |
| 4 | | + | 0412 | 0 | 0 | 0 | 1 | 1 | 1 | 1 | 1 | 1 | 1 |
| 5 | ESBL | - | 0514 | 0 | 1 | 1 | 1 | 1 | 1 | 1 | 1 | 1 | 1 |
| 5 | ESBL | - | 0510 | 0 | 0 | 1 | 0 | 1 | 1 | 1 | 1 | 1 | 1 |
| 5 | ESBL | - | 0511 | 0 | 0 | 1 | 1 | 1 | 1 | 1 | 1 | 1 | 1 |
| 5 | ESBL | - | 0513 | 0 | 0 | 0 | 0 | 1 | 1 | 1 | 1 | 1 | 1 |
| 5 | ESBL | - | 0505 | 0 | 0 | 0 | D | D | D | D | D | D | D |
| 5 | | - | 0501 | 0 | 0 | 0 | 0 | 1 | 1 | 1 | 1 | 1 | 1 |
| 5 | | - | 0512 | 0 | 0 | 0 | 0 | 1 | 1 | 1 | 1 | 1 | 1 |
| 5 | | - | 0504 | 0 | 0 | 0 | 0 | 0 | 1 | 1 | 1 | 1 | 1 |
| 5 | | - | 0507 | 0 | 0 | 0 | 0 | 0 | 1 | 1 | 1 | 1 | 1 |
| 5 | | - | 0503 | 0 | 0 | 0 | D | D | D | D | D | D | D |
| 6 | ESBL | - | 0606 | 0 | 0 | 1 | 1 | 1 | 1 | 1 | 1 | 1 | 1 |
| 6 | ESBL | - | 0612 | 0 | 0 | 1 | 1 | 1 | 1 | 1 | 1 | 1 | 1 |
| 6 | ESBL | - | 0602 | 0 | 0 | 0 | 1 | 1 | 1 | 1 | 1 | 1 | 1 |
| 6 | ESBL | - | 0614 | 0 | 0 | 0 | 0 | 1 | 1 | 1 | 1 | 1 | 1 |
| 6 | ESBL | - | 0610 | 0 | 0 | 0 | 0 | 0 | 1 | 1 | 1 | 1 | 1 |
| 6 | | - | 0605 | 0 | 0 | 0 | 0 | 1 | 0 | 1 | 1 | 1 | 1 |
| 6 | | - | 0611 | 0 | 0 | 0 | 0 | 1 | 1 | 1 | 1 | 1 | 1 |
| 6 | | - | 0615 | 0 | 0 | 0 | 0 | 1 | 1 | 1 | 1 | 1 | 1 |
| 6 | | - | 0604 | 0 | 0 | 0 | 0 | 0 | 1 | 1 | 1 | 1 | 1 |
| 6 | | - | 0613 | 0 | 0 | 0 | 0 | 0 | 1 | 1 | 1 | 1 | 1 |
| 7 | ESBL | + | 0701 | 0 | 1 | 1 | 1 | 1 | 1 | 1 | 1 | 1 | 1 |
| 7 | ESBL | + | 0702 | 0 | 1 | 1 | 1 | 1 | 1 | 1 | 1 | 1 | 1 |
| 7 | ESBL | + | 0705 | 0 | 0 | 1 | 1 | 1 | 1 | 1 | 1 | 1 | 1 |
| 7 | ESBL | + | 0708 | 0 | 0 | 1 | 1 | 1 | 1 | 1 | 1 | 1 | 1 |
| 7 | ESBL | + | 0713 | 0 | 0 | 1 | 1 | 1 | 1 | 1 | 1 | 1 | 1 |
| 7 | | + | 0703 | 0 | 0 | 1 | 1 | 1 | 1 | 1 | 1 | 1 | 1 |
| 7 | | + | 0709 | 0 | 0 | 1 | 1 | 1 | 1 | 1 | 1 | 1 | 1 |
| 7 | | + | 0710 | 0 | 0 | 1 | 1 | 1 | 1 | 1 | 1 | 1 | 1 |

| | | | | | | | | | | | | | |
|---|---|---|---|---|---|---|---|---|---|---|---|---|---|
| 7 | | + | 0712 | 0 | 0 | 1 | 1 | 1 | 1 | 1 | 1 | 1 | 1 |
| 7 | | + | 0704 | 0 | 0 | 1 | 1 | 1 | 1 | 1 | D | D | D |
| 8 | ESBL | + | 0807 | 0 | 1 | 1 | 1 | 1 | 1 | 1 | 1 | 1 | 1 |
| 8 | ESBL | + | 0804 | 0 | 0 | 1 | 1 | 1 | 1 | 1 | 1 | 1 | 1 |
| 8 | ESBL | + | 0810 | 0 | 0 | 1 | 1 | 1 | 1 | 1 | 1 | 1 | 1 |
| 8 | ESBL | + | 0812 | 0 | 0 | 1 | 1 | 1 | 1 | 1 | 1 | 1 | 1 |
| 8 | ESBL | + | 0811 | 0 | 0 | 1 | D | D | D | D | D | D | D |
| 8 | | + | 0801 | 0 | 0 | 1 | 1 | 1 | 1 | 1 | 1 | 1 | 1 |
| 8 | | + | 0802 | 0 | 0 | 1 | 1 | 1 | 1 | 1 | 1 | 1 | 1 |
| 8 | | + | 0805 | 0 | 0 | 1 | 1 | 1 | 1 | 1 | 1 | 1 | 1 |
| 8 | | + | 0809 | 0 | 0 | 1 | 1 | 1 | 1 | 1 | 1 | 1 | 1 |
| 8 | | + | 0815 | 0 | 0 | 1 | 1 | 1 | 1 | 1 | 1 | 1 | 1 |
| 9 | catA1 | - | 0906 | 0 | 0 | 1 | 1 | 0 | 1 | 1 | 1 | 1 | 1 |
| 9 | catA1 | - | 0912 | 0 | 0 | 1 | 1 | 0 | 1 | 1 | 1 | 1 | 1 |
| 9 | catA1 | - | 0915 | 0 | 0 | 1 | 0 | 1 | 1 | 1 | 1 | 1 | 1 |
| 9 | catA1 | - | 0908 | 0 | 0 | 0 | 1 | 0 | 1 | 1 | 1 | 1 | 1 |
| 9 | catA1 | - | 0913 | 0 | 0 | 0 | 1 | 1 | 1 | 1 | 1 | 1 | 1 |
| 9 | | - | 0901 | 0 | 0 | 0 | 1 | 1 | 1 | 1 | 1 | 1 | 1 |
| 9 | | - | 0902 | 0 | 0 | 0 | 1 | 1 | 1 | 1 | 1 | 1 | 1 |
| 9 | | - | 0910 | 0 | 0 | 0 | 1 | 0 | 1 | 1 | 1 | 1 | 1 |
| 9 | | - | 0914 | 0 | 0 | 0 | 1 | 1 | 1 | 1 | 1 | 1 | 1 |
| 9 | | - | 0904 | 0 | 0 | 0 | 1 | 0 | D | D | D | D | D |
| 10 | catA1 | - | 1002 | 0 | 1 | 0 | 1 | 0 | 0 | 0 | 1 | 1 | 1 |
| 10 | catA1 | - | 1004 | 0 | 1 | 0 | 0 | 0 | 0 | 0 | 1 | 1 | 1 |
| 10 | catA1 | - | 1010 | 0 | 1 | 0 | 0 | 0 | 0 | 0 | 1 | 0 | 1 |
| 10 | catA1 | - | 1013 | 0 | 1 | 0 | 0 | 0 | 0 | 0 | 1 | 1 | 1 |
| 10 | catA1 | - | 1003 | 0 | 0 | 0 | 1 | 0 | 0 | 0 | 1 | 1 | 1 |
| 10 | | - | 1006 | 0 | 0 | 0 | 1 | 0 | 0 | 0 | 1 | 1 | 1 |
| 10 | | - | 1007 | 0 | 0 | 0 | 1 | 0 | 0 | 0 | 1 | 1 | 1 |
| 10 | | - | 1009 | 0 | 0 | 0 | 1 | 0 | 0 | 1 | 1 | 1 | 1 |
| 10 | | - | 1005 | 0 | 0 | 0 | 0 | 0 | 0 | 1 | 1 | 1 | 1 |
| 10 | | - | 1015 | 0 | 0 | 0 | 0 | 0 | 0 | 0 | 1 | 1 | 1 |
| 11 | catA1 | + | 1103 | 0 | 0 | 0 | 0 | 0 | 1 | 1 | 1 | 1 | 1 |
| 11 | catA1 | + | 1104 | 0 | 0 | 0 | 0 | 0 | 1 | 1 | 1 | 1 | 1 |
| 11 | catA1 | + | 1105 | 0 | 0 | 0 | 0 | 0 | 1 | 1 | 1 | 1 | 1 |
| 11 | catA1 | + | 1110 | 0 | 0 | 0 | 0 | 0 | 1 | 1 | 1 | 1 | 1 |
| 11 | catA1 | + | 1112 | 0 | 0 | 0 | 0 | 0 | 1 | 1 | 1 | 1 | 1 |
| 11 | | + | 1102 | 0 | 0 | 0 | 0 | 0 | 1 | 1 | 1 | 1 | 1 |
| 11 | | + | 1106 | 0 | 0 | 0 | 0 | 0 | 1 | 1 | 1 | 1 | 1 |
| 11 | | + | 1107 | 0 | 0 | 0 | 0 | 0 | 1 | 1 | 1 | 1 | 1 |
| 11 | | + | 1108 | 0 | 0 | 0 | 0 | 0 | 1 | 1 | 1 | 1 | 1 |
| 11 | | + | 1115 | 0 | 0 | 0 | 0 | 0 | 1 | 1 | 1 | 1 | 1 |
| 12 | catA1 | + | 1204 | 0 | 0 | 1 | 1 | 1 | 1 | 1 | 1 | 1 | 1 |
| 12 | catA1 | + | 1207 | 0 | 0 | 1 | 1 | 1 | 1 | 1 | 1 | 1 | 1 |
| 12 | catA1 | + | 1209 | 0 | 0 | 1 | 1 | 1 | 1 | 1 | 1 | 1 | 1 |
| 12 | catA1 | + | 1214 | 0 | 0 | 0 | 1 | 1 | 1 | 1 | 1 | 1 | 1 |

| | | | | | | | | | | | | | |
|---|---|---|---|---|---|---|---|---|---|---|---|---|---|
| 12 | catA1 | + | 1213 | 0 | 0 | 0 | 0 | 1 | 1 | 1 | 1 | 1 | 1 |
| 12 | | + | 1201 | 0 | 0 | 1 | 1 | 1 | 1 | 1 | 1 | 1 | 1 |
| 12 | | + | 1203 | 0 | 0 | 1 | 1 | 1 | 1 | 1 | 1 | 1 | 1 |
| 12 | | + | 1210 | 0 | 0 | 1 | 1 | 1 | 1 | 1 | 1 | 1 | 1 |
| 12 | | + | 1202 | 0 | 0 | 0 | 1 | 1 | 1 | 1 | 1 | 1 | 1 |
| 12 | | + | 1212 | 0 | 0 | 0 | 0 | 1 | 1 | 1 | 1 | 1 | 1 |

##### 1.2. Protocols to adjust raw transmission data

Since all broilers were excreting (status “1” in Table S1) at the end of the experiment, we assumed transmission of *E. coli* bacteria between broilers follows SI-dynamics in which broilers will remain excreting (status “1” in Table S1) until the end of the experiment once their cloacal swabs test positive for the resistant bacteria. However, several broilers with a positive sample became negative at the following sampling time point, and then positive again at a later sampling time point. We adjusted the status of these negative samples according to 1 of the 3 protocols described below and fitted a Bayesian hierarchical model to each of these adjusted data sets. Model diagnostics indicated the models fitted all 3 data sets well. We selected the third protocol to create our final input data because considering reoccurrences of negative samples to be false negatives best reflects the nature of transmission and sequencing, with false-negative samples arising because of limitations in sampling and detecting small numbers of resistant bacteria.

###### 1.2.1. Protocol 1: Only a single reoccurrence of negative status is treated as a false negative

Rules to change the data in each row from the first occurrence of a positive swab (i.e., status “1”):

- 1: When “1” is followed by a single “0”, this “0” is assumed to be a false negative and is changed to “1”.
- 2: When “1” is followed by more than one “0”, all preceding “1” are assumed to be false positive and are changed to “0”.
- 3: When “1” is followed by “0” followed by “D” (i.e., the broiler died), the “0” is assumed to be a false negative and is changed to “1”.

###### 1.2.2. Protocol 2: Prolonged reoccurrence of a negative status is treated as false negative if it follows a single occurrence of positive status

Rules to change the data in each row from the first occurrence of a positive swab (i.e., status “1”):

- 1: When “1” is followed by a single “0”, this “0” is assumed to be a false negative and is changed to “1”.
- 2: When a single “1” is followed by more than one “0”, this “1” is assumed to be a false positive and is changed to “0”.
- 3: When more than one “1” is followed by any number of “0”, the “0” are assumed to be false negative and are changed to “1”
- 4: When “1” is followed by “0” followed by “D” (i.e., the broiler died), the “0” is assumed to be a false negative and is changed to “1”.

1.2.3. <u>Protocol 3: Each reoccurrence of a negative status is treated as a false negative</u>

Rules to change the data in each row from the first occurrence of a positive swab (i.e., status “1”):

- 1: When “1” is followed by any number of “0”, all “0” are assumed to be false negative and are changed to “1”.

*Table S2: Examples of the status of 3 broilers at different time points (hours after inoculation) giving the raw data and adjustments according to each of the 3 protocols. The last time point recorded status 1 for all entries and has been omitted to reduce the width of the table. Shading indicates changes compared to the raw data. Abbreviations: ID: broiler ID, P: protocol*

| ID | 0113 | | | | | | | | | 1002 | | | | | | | | | 1004 | | | | | | | | |
|---|---|---|---|---|---|---|---|---|---|---|---|---|---|---|---|---|---|---|---|---|---|---|---|---|---|---|---|
| time | 0 | 8 | 24 | 32 | 48 | 72 | 96 | 120 | 168 | 0 | 8 | 24 | 32 | 48 | 72 | 96 | 120 | 168 | 0 | 8 | 24 | 32 | 48 | 72 | 96 | 120 | 168 |
| raw | 0 | 0 | 0 | 0 | 0 | 1 | 0 | 1 | 1 | 0 | 1 | 0 | 1 | 0 | 0 | 0 | 1 | 1 | 0 | 1 | 0 | 0 | 0 | 0 | 0 | 1 | 1 |
| P1 | 0 | 0 | 0 | 0 | 0 | 1 | 1 | 1 | 1 | 0 | 0 | 0 | 0 | 0 | 0 | 0 | 1 | 1 | 0 | 0 | 0 | 0 | 0 | 0 | 0 | 1 | 1 |
| P2 | 0 | 0 | 0 | 0 | 0 | 1 | 1 | 1 | 1 | 0 | 1 | 1 | 1 | 1 | 1 | 1 | 1 | 1 | 0 | 0 | 0 | 0 | 0 | 0 | 0 | 1 | 1 |
| P3 | 0 | 0 | 0 | 0 | 0 | 1 | 1 | 1 | 1 | 0 | 1 | 1 | 1 | 1 | 1 | 1 | 1 | 1 | 0 | 1 | 1 | 1 | 1 | 1 | 1 | 1 | 1 |

##### 1.3. Distribution of sexes

*Table S3: Overview of the distribution of the sexes in the different experimental groups. Abbreviations: F: female, M: male, No: pens without antibiotics, T: total, Yes: pens with antibiotics.*

| | | Day 5 | | | | Day 14 | | | | Total | | | |
|---|---|---|---|---|---|---|---|---|---|---|---|---|---|
| Group | Pens | F | M | T | %F | F | M | T | %F | F | M | T | %F |
| CPE_No | 1, 2 | 6 | 4 | 10 | 60 | 9 | 7 | 16 | 56 | 15 | 11 | 26 | 58 |
| ESBL_No | 5, 6 | 8 | 2 | 10 | 80 | 8 | 10 | 18 | 44 | 16 | 12 | 28 | 57 |
| catA1_No | 9, 10 | 6 | 4 | 10 | 60 | 8 | 11 | 19 | 42 | 14 | 15 | 29 | 48 |
| CPE_Yes | 3, 4 | 4 | 5 | 9 | 44 | 10 | 10 | 20 | 50 | 14 | 15 | 29 | 48 |
| ESBL_Yes | 7, 8 | 6 | 4 | 10 | 60 | 11.5 | 6.5 | 18 | 64 | 17.5 | 10.5 | 28 | 63 |
| catA1_Yes | 11, 12 | 2 | 7 | 9 | 22 | 10 | 10 | 20 | 50 | 12 | 17 | 29 | 41 |
| CPE | 1 - 4 | 10 | 9 | 19 | 53 | 19 | 17 | 36 | 53 | 29 | 26 | 55 | 53 |
| ESBL | 5 - 8 | 14 | 6 | 20 | 70 | 19.5 | 16.5 | 36 | 54 | 33.5 | 22.5 | 56 | 60 |
| catA1 | 9 - 12 | 8 | 11 | 19 | 42 | 18 | 21 | 39 | 46 | 26 | 32 | 58 | 45 |
| No | 1, 2, 5, 6, 9, 10 | 20 | 10 | 30 | 67 | 25 | 28 | 53 | 47 | 45 | 38 | 83 | 54 |
| Yes | 3, 4, 7, 8, 11, 12 | 12 | 16 | 28 | 43 | 31.5 | 26.5 | 58 | 54 | 43.5 | 42.5 | 86 | 51 |
| Total | 1 - 12 | 32 | 26 | 58 | 55 | 56.5 | 54.5 | 111 | 51 | 88.5 | 80.5 | 169 | 52 |

### 2. Data from the microbiome analysis

##### 2.1. Quality controls for microbiome sequencing

2.1.1. <u>Composition of the spiked negative controls</u>

The composition of the negative controls spiked with a low concentration of microbial community DNA standard (ZymoBIOMICS; Zymo Research Corporation, Irvine, CA) was similar for all controls, was close to the theoretical composition indicated by the manufacturer, and contained less than 0.15% foreign DNA (Figure S1). This shows the sequencing runs went well, the use of multiple 96-well plates did not lead to large deviations in obtained composition, and little contamination occurred.

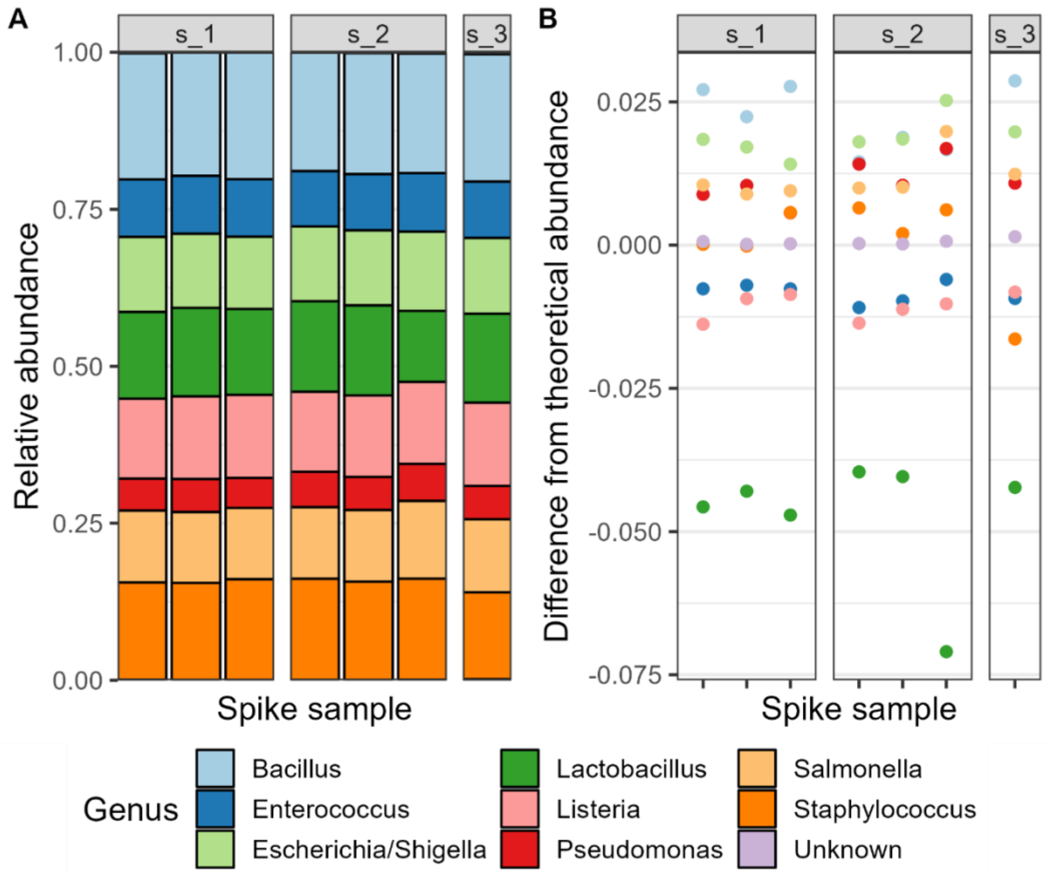


*Figure S1: A: composition of the spike samples at genus level as determined by sequencing. B: The difference between the observed and theoretical composition at genus level. s_1, s_2, and s_3 indicate the different well plates the spike samples were on.*

2.1.2. Number of reads

Slightly less than half of all sequences could not be assigned on genus level after removing the spike samples and non-bacterial sequences, with better coverage on higher taxonomic levels (Table S4). We left unassigned sequences as-is, i.e., we did not use the subsequent higher taxonomic ranks to fill in NAs.

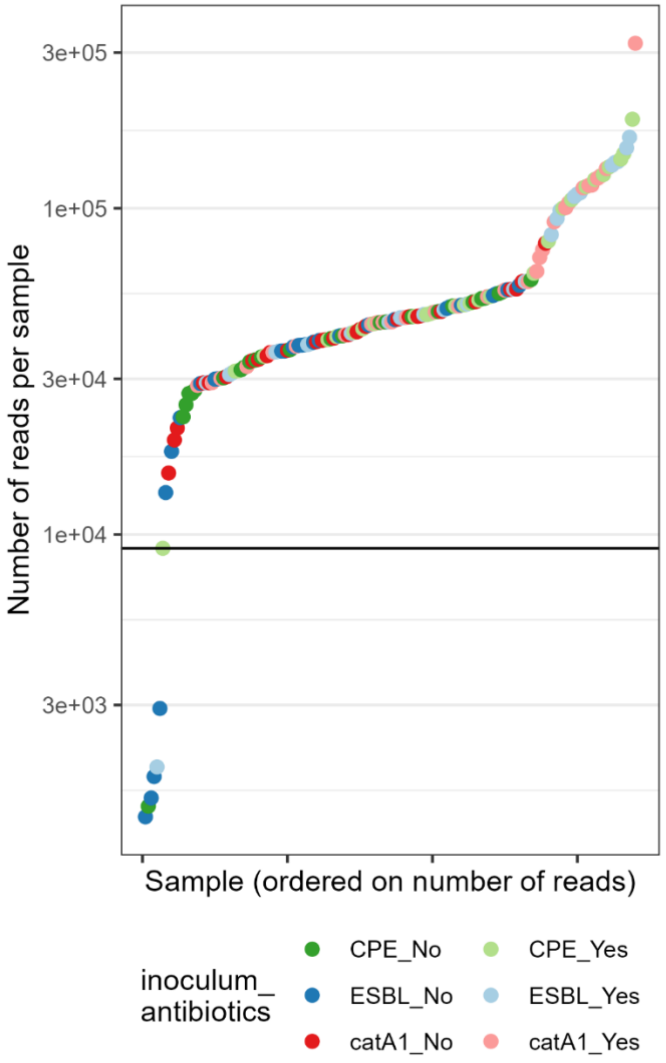


Figure S2: The number of reads in the samples. The black horizontal line indicates the number of 9071 reads in the sample with the 7th least number of reads. Colours indicate the absence ('No') or presence ('Yes') of antibiotic treatment and the different inoculums (CPE-strain, ESBL-strain, catA1-strain).

*Table S4: Numbers and percentages of sequences not assigned to a taxon for each taxonomic rank, after removal of the non-bacterial sequences and the spiked negative controls.*

| Rank | n NA | % NA |
|---|---|---|
| Phylum | 53 | 0.7 |
| Class | 152 | 1.9 |
| Order | 608 | 7.7 |
| Family | 1406 | 17.7 |
| Genus | 3715 | 46.8 |

2.1.3. Rarefaction curves

Rarefaction curves on amplicon sequence variant (ASV) level for samples without antibiotics on day 5, and for most samples on day 14 did not level off (bottom row in Figure S3), indicating not all ASVs present in the sample have been sequenced. This is less prominent at the genus level (top row in Figure S3).

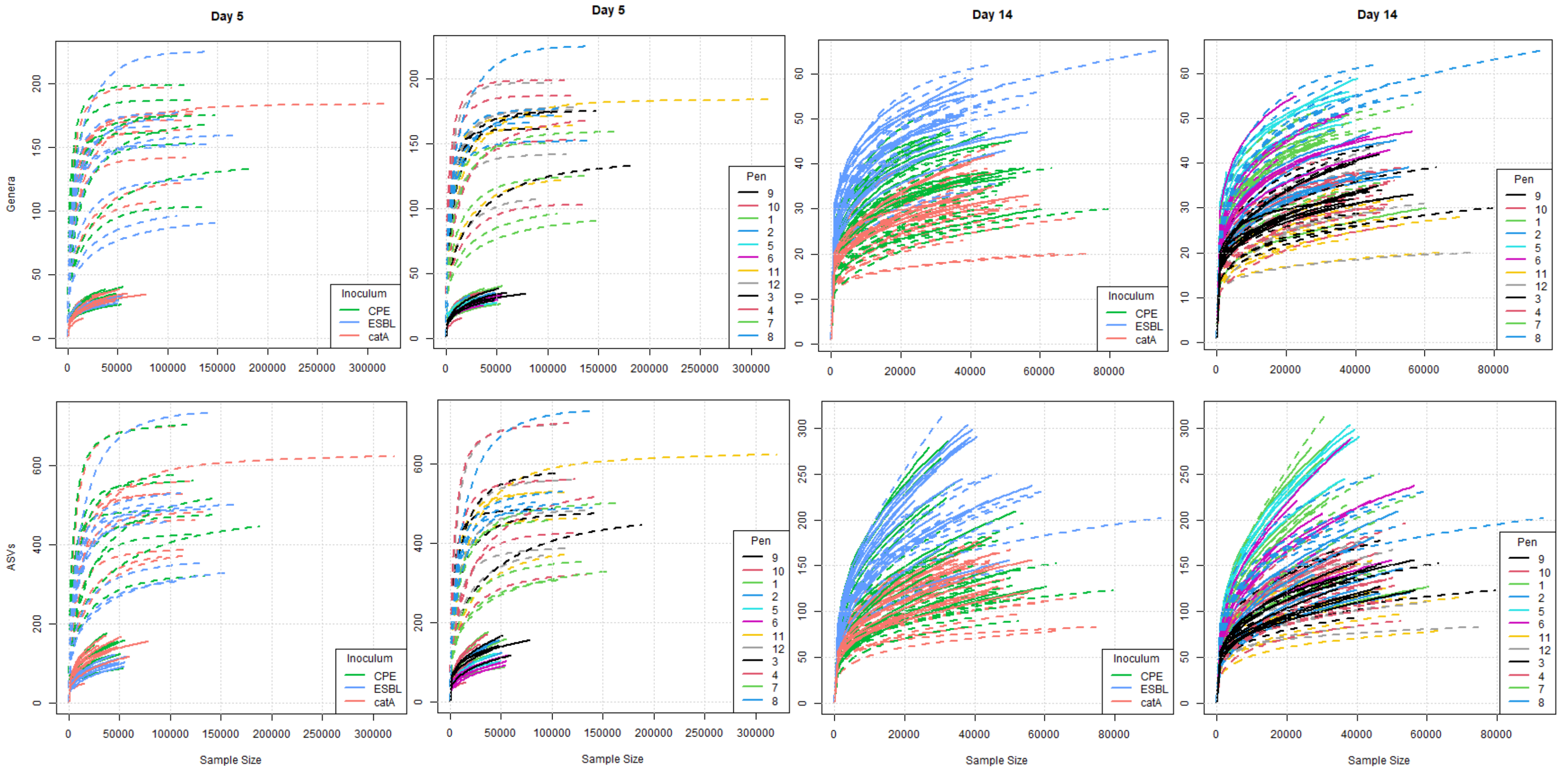


*Figure S3: Rarefaction curves for samples on day 5 (first 2 columns) and day 14 (last 2 columns) at genus level (top row) and ASV level (bottom row). Solid lines indicate samples without antibiotic treatment and dashed lines indicate samples with antibiotic treatment. Note the axes differ between days and between the genus level and ASV level.*

##### 2.2. Microbiome composition

The microbiome of all broilers was dominated by the classes Gammaproteobacteria, Clostridia, and Bacilli (
Figure S4). On day 5 the distinction between groups with and without antibiotics is clear at class, family and genus level (
Figure S4; Figure S5; Figure S6). On day 14, the composition in pens 11 and 12 (catA1-strain with antibiotics) resembles the composition in pens 9 and 10 (catA1-strain without antibiotics) more closely than the composition in other pens with antibiotics at family level (Figure S5), which could reflect a room effect as pens 9 to 12 were in the same room. This is not the case at class level (
Figure S4).

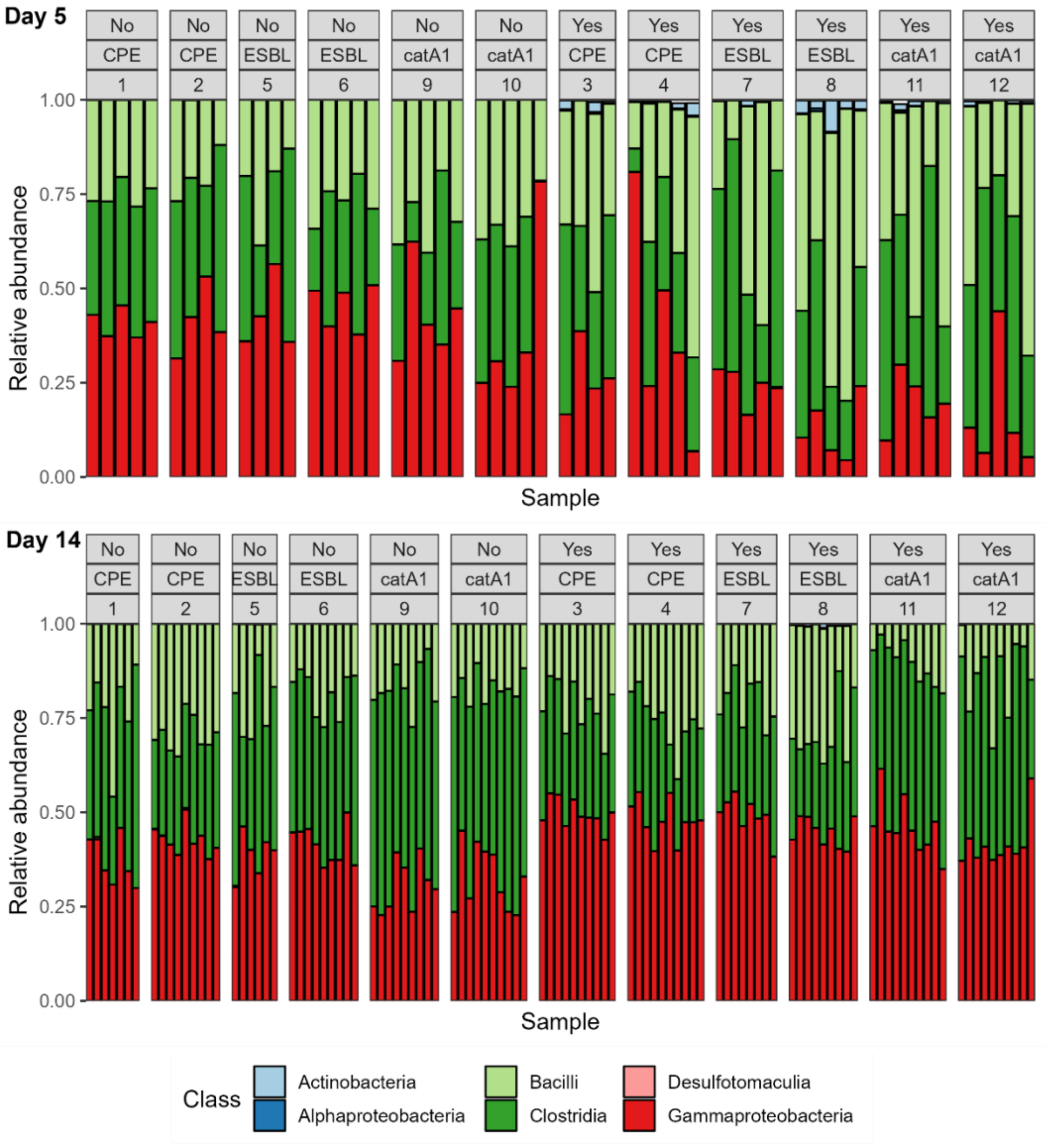


Figure S4: Relative abundance (vertical axis) of the 6 most-abundant classes considering the total abundance over all samples. Facet labels indicate the absence ('No') or presence ('Yes') of antibiotic treatment, the different inoculums (CPE-strain, ESBL-strain, catA1-strain), and the pen number.

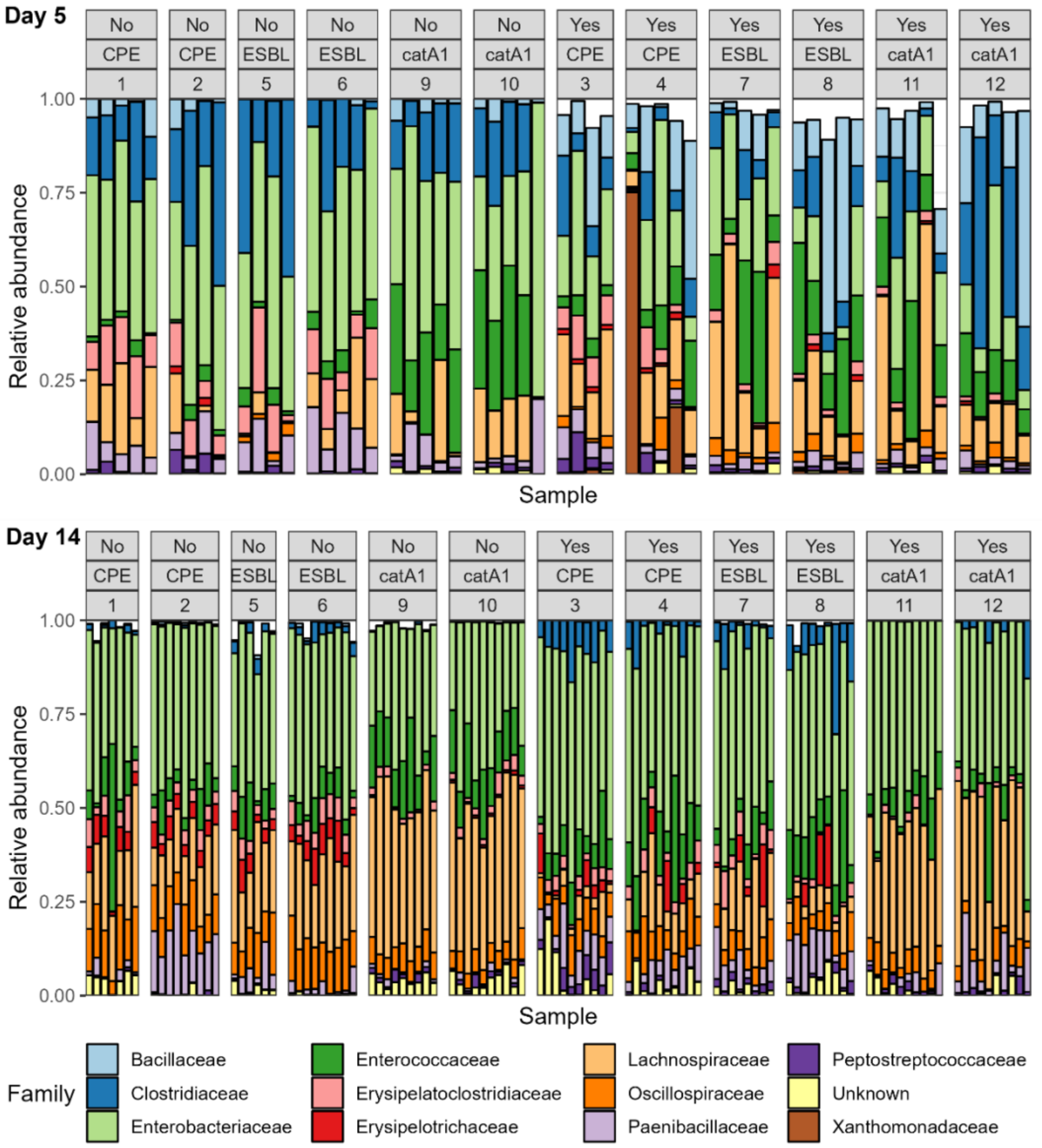


*Figure S5: Relative abundance (vertical axis) of the 12 most-abundant families considering the total abundance over all samples. Facet labels indicate the absence ('No') or presence ('Yes') of antibiotic treatment, the different inoculums (CPE-strain, ESBL-strain, catA1-strain), and the pen number.*

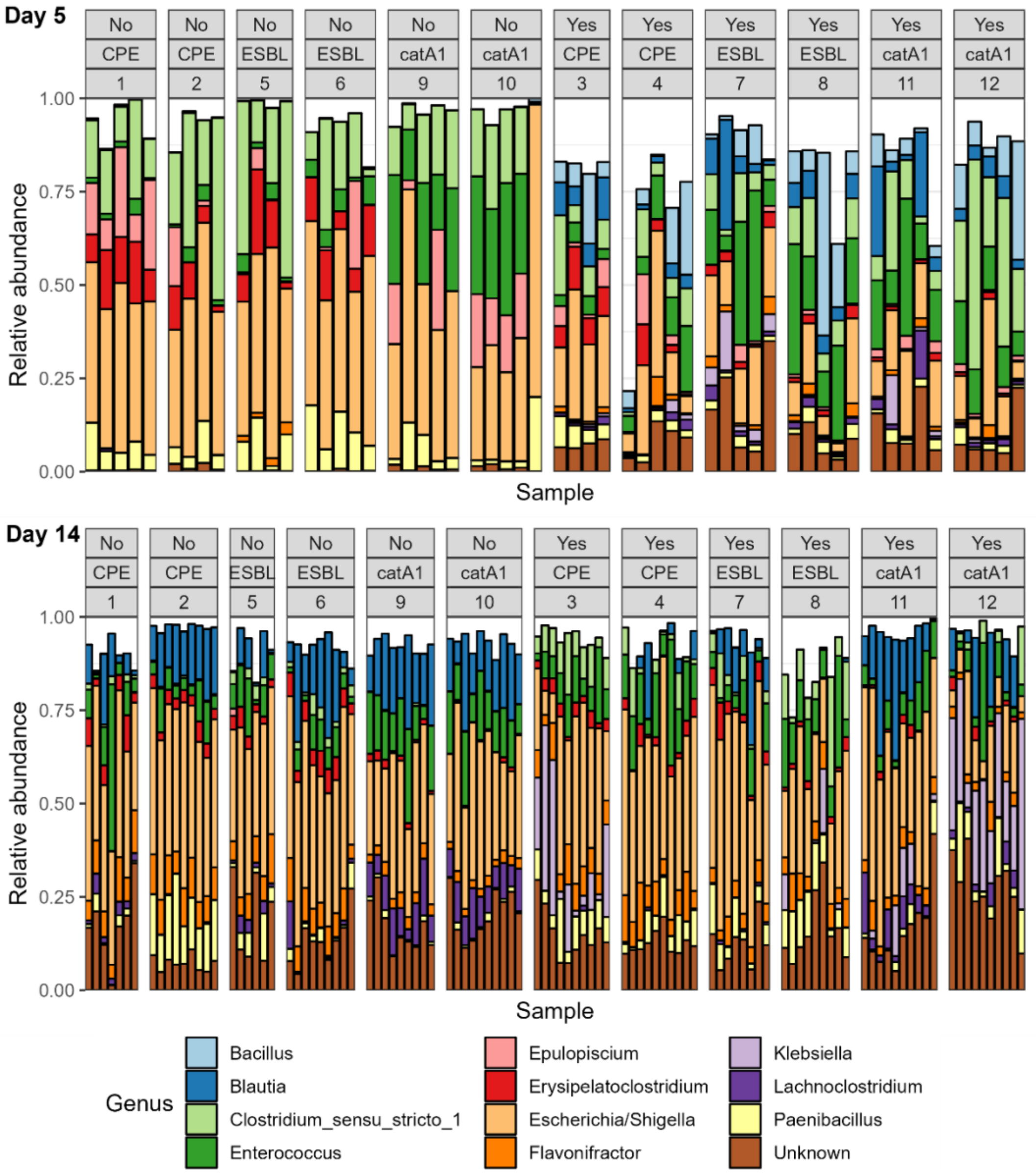


*Figure S6: Relative abundance (vertical axis) of the 12 most-abundant genera considering the total abundance over all samples. Facet labels indicate the absence ('No') or presence ('Yes') of antibiotic treatment, the different inoculums (CPE-strain, ESBL-strain, catA1-strain), and the pen number.*

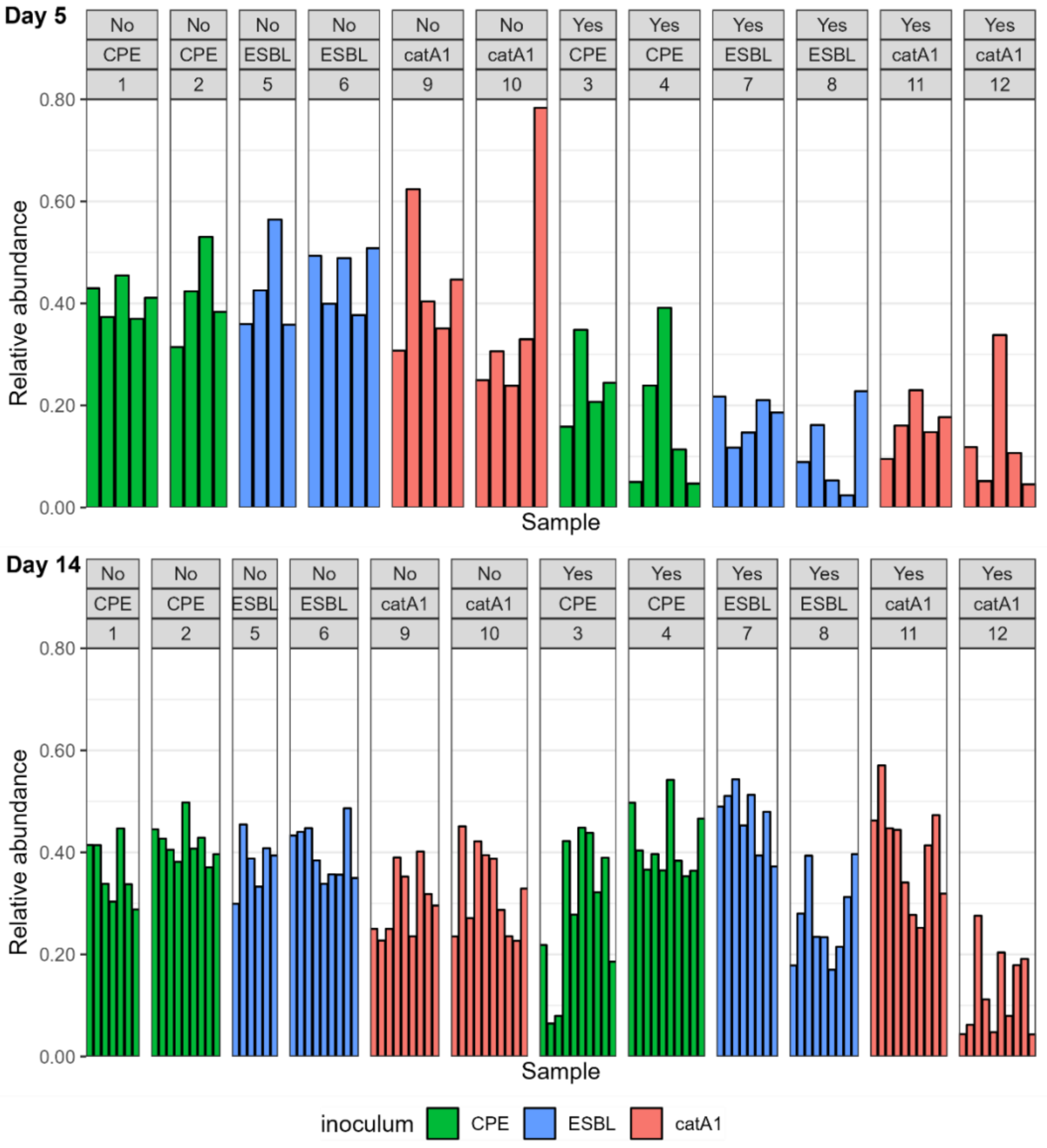


*Figure S7:* Relative abundance (vertical axis) of the *Escherichia/Shigella genus*. Facet labels indicate the absence ('No') or presence ('Yes') of antibiotic treatment, the different inoculums (CPE-strain, ESBL-strain, catA1-strain), and the pen number.

#### 2.3. Alpha-diversity on ASV level

As expected, because caecal samples on day 5 were taken before inoculation, observed richness, Shannon's diversity, and Pielou's evenness at ASV level on day 5 were not different between groups inoculated with the different inoculums (i.e., CPE-strain, ESBL-strain, catA1-strain; Figure S8 and Table S7). Observed richness on day 14 was slightly higher in broilers inoculated with the ESBL-strain than in broilers inoculated with the CPE-strain or catA1-strain, both in the non-amoxicillin-treated group and in the amoxicillin-treated group (Figure S8 and Table S8). Shannon's diversity on day 14 was not different between the inoculums. Pielou's evenness on day 14 was higher in the catA1-strain than in the CPE-strain or ESBL-strain in groups without and with antibiotics.

Observed richness and Shannon's diversity at ASV level on day 5 were lower in the non-amoxicillin-treated groups than in the amoxicillin-treated groups, but Pielou's evenness was not different (Figure S8 and

Table S7) indicating fewer different ASV were present in the non-amoxicillin-treated groups but the distribution of their abundances was similar to the distribution of their abundance in the amoxicillin-treated groups. Shannon's diversity on day 14 was higher in non-amoxicillin-treated groups than in amoxicillin-treated groups, but no differences in observed richness or Pielou's evenness were found between those groups (Figure S8 and

Table S8).

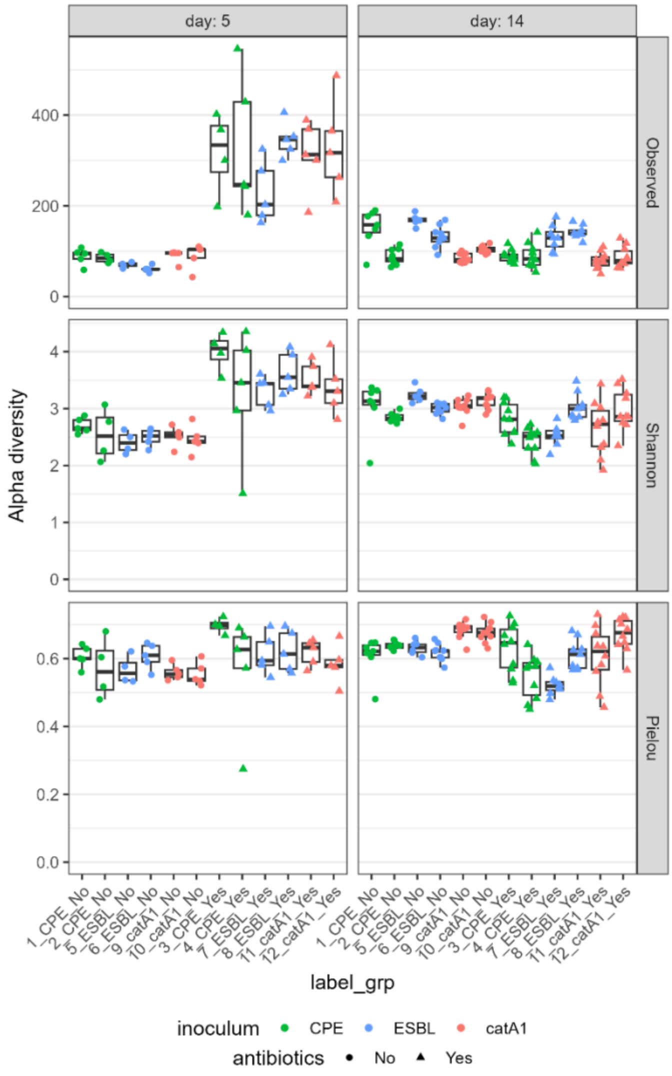


*Figure S8: Alpha-diversity (y-axis) by inoculum and antibiotic treatment (horizontal axis) at ASV level. Colours indicate different inoculums (CPE-strain: green; ESBL-strain: blue; catA1-strain: red) and symbols indicate the absence (circles) or presence (triangles) of antibiotic treatment. The panels show the different alpha-diversity measures (rows) and different days (columns).*

#### 2.4. Alpha-diversity: tables

*Table S5: Differences in alpha-diversity between groups at genus level on day 5 based on Kruskal-Wallis rank sum test and post hoc Dunn's test. P-values have been adjusted with Benjamini-Hochberg correction. 'No' and 'Yes' refers to pens without and with antibiotics, respectively.*

| | Observed | | Shannon | | Pielou | |
|---|---|---|---|---|---|---|
| comparisons | Z | P.adj | Z | P.adj | Z | P.adj |
| CPE_No – ESBL_No | 0.782 | 0.6516 | 0.795 | 0.6397 | 0.795 | 0.5816 |
| CPE_No – catA1_No | 0.401 | 0.8605 | 0.523 | 0.8195 | 1.077 | 0.4694 |
| ESBL_No – catA1_No | -0.401 | 0.9388 | -0.293 | 0.8246 | 0.261 | 0.8510 |
| CPE_Yes – ESBL_Yes | 0.272 | 0.8418 | 0.181 | 0.8566 | 1.186 | 0.4418 |
| CPE_Yes – catA1_Yes | 0.003 | 0.9977 | 0.469 | 0.7987 | 1.789 | 0.2208 |
| ESBL_Yes – catA1_Yes | -0.276 | 0.9026 | 0.296 | 0.8849 | 0.620 | 0.6178 |
| CPE_No – CPE_Yes | -3.347 | 0.0015 | -3.181 | 0.0031 | -1.832 | 0.3349 |
| ESBL_No – ESBL_Yes | -3.964 | 0.0004 | -3.899 | 0.0007 | -1.509 | 0.3280 |
| catA1_No – catA1_Yes | -3.937 | 0.0003 | -3.408 | 0.0016 | -1.199 | 0.4940 |

*Table S6: Differences in alpha-diversity between groups at genus level on day 14 based on Kruskal-Wallis rank sum test and post hoc Dunn's test. P-values have been adjusted with Benjamini-Hochberg correction. 'No' and 'Yes' refers to pens without and with antibiotics, respectively.*

| | Observed | | Shannon | | Pielou | |
|---|---|---|---|---|---|---|
| comparisons | Z | P.adj | Z | P.adj | Z | P.adj |
| CPE_No – ESBL_No | -2.192 | 0.0425 | -1.602 | 0.4094 | 0.145 | 1.0000 |
| CPE_No – catA1_No | 2.295 | 0.0362 | -1.190 | 0.3903 | -3.319 | 0.0045 |
| ESBL_No – catA1_No | 4.536 | 0.0000 | 0.498 | 0.7729 | -3.411 | 0.0048 |
| CPE_Yes – ESBL_Yes | -4.522 | 0.0000 | -1.286 | 0.4250 | 2.322 | 0.0759 |
| CPE_Yes – catA1_Yes | 1.446 | 0.1709 | 1.269 | 0.3835 | 0.510 | 0.7630 |
| ESBL_Yes – catA1_Yes | 5.908 | 0.0000 | 2.502 | 0.0925 | -1.834 | 0.1430 |
| CPE_No – CPE_Yes | 2.566 | 0.0193 | -0.091 | 0.9273 | -2.172 | 0.0745 |
| ESBL_No – ESBL_Yes | 0.442 | 0.7051 | 0.341 | 0.8459 | -0.042 | 0.9664 |
| catA1_No – catA1_Yes | 1.683 | 0.1155 | 2.417 | 0.0783 | 1.743 | 0.1355 |

*Table S7: Differences in alpha-diversity between groups at ASV level on day 5 based on Kruskal-Wallis rank sum test and post hoc Dunn's test. P-values have been adjusted with Benjamini-Hochberg correction. 'No' and 'Yes' refers to pens without and with antibiotics, respectively.*

| | Observed | | Shannon | | Pielou | |
|---|---|---|---|---|---|---|
| comparisons | Z | P.adj | Z | P.adj | Z | P.adj |
| CPE_No – ESBL_No | 1.214 | 0.3063 | 0.88 | 0.5679 | 0.213 | 0.8313 |
| CPE_No – catA1_No | -0.224 | 0.8818 | 0.801 | 0.5767 | 1.584 | 0.2832 |
| ESBL_No – catA1_No | -1.470 | 0.2125 | -0.102 | 0.9188 | 1.365 | 0.3229 |
| CPE_Yes – ESBL_Yes | 0.310 | 0.8733 | 0.255 | 0.9216 | 1.072 | 0.4726 |
| CPE_Yes – catA1_Yes | -0.077 | 0.9384 | 0.360 | 0.8987 | 1.557 | 0.2558 |
| ESBL_Yes – catA1_Yes | -0.397 | 0.8638 | 0.108 | 0.9795 | 0.498 | 0.7727 |
| CPE_No – CPE_Yes | -3.409 | 0.0016 | -2.982 | 0.0061 | -1.789 | 0.2207 |
| ESBL_No – ESBL_Yes | -4.433 | 0.0000 | -3.708 | 0.0010 | -0.982 | 0.4892 |
| catA1_No – catA1_Yes | -3.443 | 0.0017 | -3.597 | 0.0008 | -1.913 | 0.2091 |

*Table S8: Differences in alpha-diversity between groups at ASV level on day 14 based on Kruskal-Wallis rank sum test and post hoc Dunn's test. P-values have been adjusted with Benjamini-Hochberg correction. 'No' and 'Yes' refers to pens without and with antibiotics, respectively.*

| | Observed | | Shannon | | Pielou | |
|---|---|---|---|---|---|---|
| comparisons | Z | P.adj | Z | P.adj | Z | P.adj |
| CPE_No – ESBL_No | -2.626 | 0.0162 | -1.522 | 0.1920 | 0.371 | 0.7109 |
| CPE_No – catA1_No | 1.121 | 0.3281 | -1.605 | 0.1808 | -3.024 | 0.0075 |
| ESBL_No – catA1_No | 3.833 | 0.0004 | 0.007 | 0.9945 | -3.356 | 0.0030 |
| CPE_Yes – ESBL_Yes | -4.292 | 0.0001 | -1.226 | 0.2751 | 1.136 | 0.3198 |
| CPE_Yes – catA1_Yes | 0.711 | 0.5112 | -1.793 | 0.1367 | -2.344 | 0.0477 |
| ESBL_Yes – catA1_Yes | 4.973 | 0.0000 | -0.493 | 0.6664 | -3.383 | 0.0036 |
| CPE_No – CPE_Yes | 1.967 | 0.0738 | 2.577 | 0.0249 | 1.177 | 0.3262 |
| ESBL_No – ESBL_Yes | 0.530 | 0.5962 | 2.842 | 0.0168 | 1.797 | 0.1207 |
| catA1_No – catA1_Yes | 1.574 | 0.1575 | 2.628 | 0.0258 | 2.122 | 0.0635 |

#### 2.5. Beta-diversity at ASV level

The inoculum explained 9% and 6% of the variation in Bray-Curtis dissimilarity and Jaccard distance at ASV level on day 5 (i.e., before inoculation), antibiotic treatment explained 22% of the variation for both measures, and their interaction explained 5% of the variation for both measures (Table S9). The different inoculums were not separate in the principal coordinate plot (Figure S9), apart from the catA1-strain without antibiotics being separate from the CPE-strain and ESBL-strain for Jaccard distance. In contrast, the groups without and with antibiotics were clearly separate. No differences in dispersion were found.

The inoculum explained 21% and 15% of the variation in Bray-Curtis dissimilarity and Jaccard distance at ASV level on day 14, antibiotic treatment explained 17% and 7% of the variation, and their interaction explained 6% and 5% of the variation (Table S10). The catA1-strains were clearly separate from the CPE-strains and ESBL-strains in the principal coordinate plots, whereas the CPE-strains and ESBL-strains overlapped much with each other (Figure S9). The groups without and with antibiotics clearly separated with Bray-Curtis dissimilarity but not with Jaccard distance, indicating they differ mostly in the most-abundant ASVs. For both beta-diversity measures, the dispersion in the catA1-strain without antibiotics was different from dispersion in all other groups, and dispersion in the CPE-strain without antibiotics was smaller than dispersion in the ESBL-strain with antibiotics and smaller than dispersion in the catA1-strain with antibiotics, such that the differences could be a difference in location, a difference in dispersion, or both.

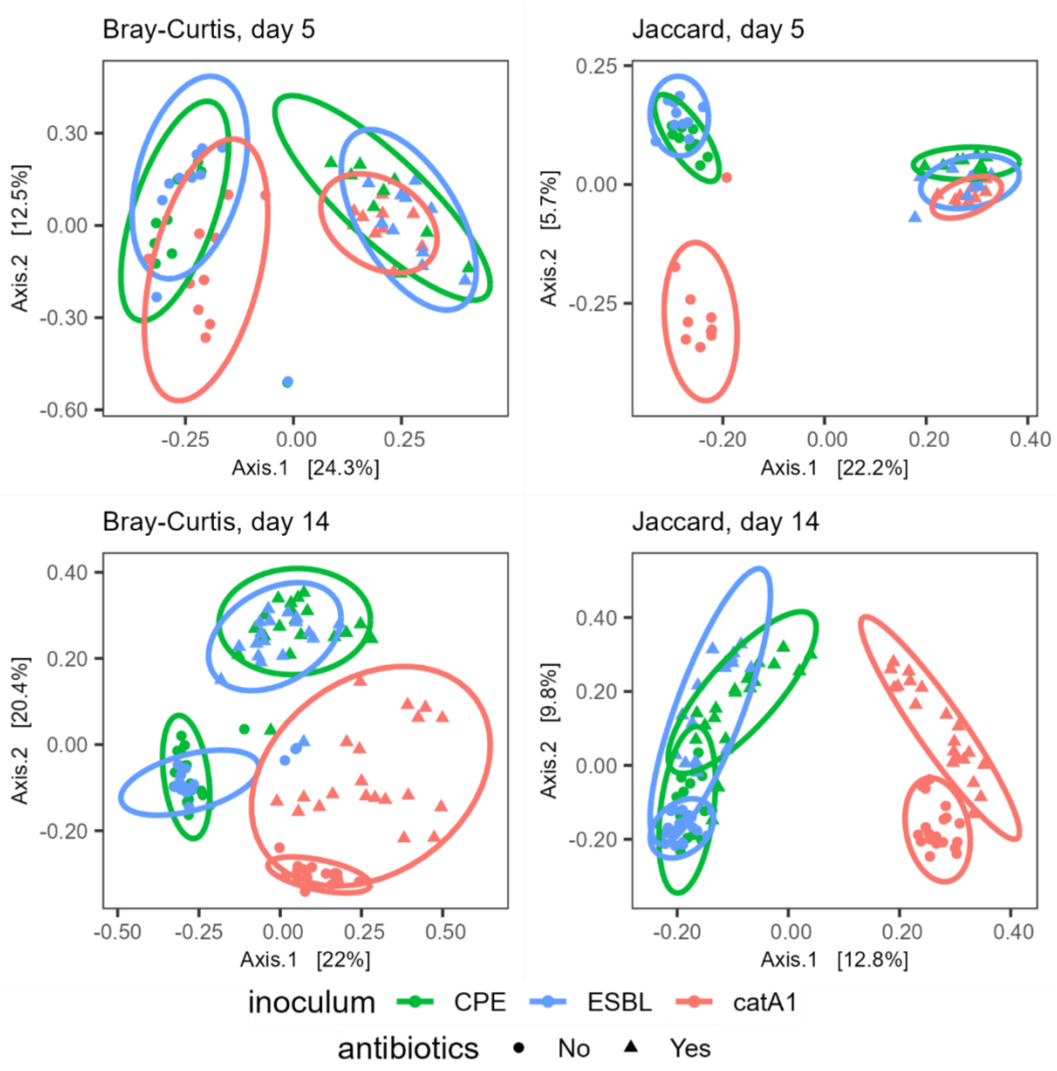


*Figure S9: Principal coordinate analysis of Bray-Curtis dissimilarity (left) and Jaccard distance (right) for day 5 (top) and day 14 (bottom) at ASV level. Colours indicate different inoculums (CPE-strain: green; ESBL-strain: blue; catA1-strain: red) and symbols indicate the absence (circles) or presence (triangles) of antibiotic treatment. Ellipses represent 95% confidence regions assuming a multivariate t-distribution.*

#### 2.6. Beta-diversity: tables

*Table S9: Permutational multivariate analysis of variance for the effects of E. coli, antibiotics, and their interaction on beta-diversity measured with Bray-Curtis dissimilarity and Jaccard distance at genus level and ASV level on day 5.*

| | Bray-Curtis dissimilarity | | | | | | Jaccard distance | | | | | |
|---|---|---|---|---|---|---|---|---|---|---|---|---|
| | Genus level | | | ASV level | | | Genus level | | | ASV level | | |
| | Df | $R^2$ | Pr(>F) | Df | $R^2$ | Pr(>F) | Df | $R^2$ | Pr(>F) | Df | $R^2$ | Pr(>F) |
| Inoculum | 2 | 0.060 | 0.003 | 2 | 0.092 | 0.001 | 2 | 0.026 | 0.140 | 2 | 0.060 | 0.001 |
| Antibiotic treatment | 1 | 0.271 | 0.001 | 1 | 0.222 | 0.001 | 1 | 0.501 | 0.001 | 1 | 0.219 | 0.001 |
| Interaction | 2 | 0.046 | 0.025 | 2 | 0.052 | 0.004 | 2 | 0.027 | 0.135 | 2 | 0.050 | 0.007 |
| Residual | 53 | 0.623 | | 53 | 0.634 | | 53 | 0.446 | | 53 | 0.672 | |
| Total | 58 | 1.000 | | 58 | 1.000 | | 58 | 1.000 | | 58 | 1.000 | |

*Table S10: Permutational multivariate analysis of variance for the effects of E. coli, antibiotics, and their interaction on beta-diversity measured with Bray-Curtis dissimilarity and Jaccard distance at genus level and ASV level on day 14.*

| | Bray-Curtis dissimilarity | | | | | | Jaccard distance | | | | | |
|---|---|---|---|---|---|---|---|---|---|---|---|---|
| | Genus level | | | ASV level | | | Genus level | | | ASV level | | |
| | Df | $R^2$ | Pr(>F) | Df | $R^2$ | Pr(>F) | Df | $R^2$ | Pr(>F) | Df | $R^2$ | Pr(>F) |
| Inoculum | 2 | 0.163 | 0.001 | 2 | 0.209 | 0.001 | 2 | 0.169 | 0.001 | 2 | 0.146 | 0.001 |
| Antibiotic treatment | 1 | 0.093 | 0.001 | 1 | 0.170 | 0.001 | 1 | 0.085 | 0.001 | 1 | 0.072 | 0.001 |
| Interaction | 2 | 0.045 | 0.002 | 2 | 0.061 | 0.001 | 2 | 0.056 | 0.001 | 2 | 0.046 | 0.001 |
| Residual | 105 | 0.700 | | 105 | 0.559 | | 105 | 0.690 | | 105 | 0.737 | |
| Total | 110 | 1.000 | | 110 | 1.000 | | 110 | 1.000 | | 110 | 1.000 | |

#### 2.7. Similarity percentage analyses: tables

*Table S11: Genera that contributed more than 1 per cent to the Bray-Curtis dissimilarity on day 5 between groups without and with antibiotic treatment in pens inoculated with CPE. 'No' and 'Yes' refers to pens without and with antibiotics, respectively. Abbreviations: p: permutation p-value.*

| Genus | Average contribution | Standard deviation | Average abundance No | Average abundance Yes | p |
|---|---|---|---|---|---|
| Escherichia/Shigella | 0.104 | 0.086 | 15239 | 27758 | 0.981 |
| Stenotrophomonas | 0.082 | 0.184 | 0 | 13163 | 0.001 |
| Bacillus | 0.072 | 0.060 | 84 | 11411 | 0.001 |
| Enterococcus | 0.050 | 0.037 | 831 | 8920 | 0.001 |
| Clostridium sensu stricto 1 | 0.047 | 0.046 | 8217 | 11052 | 1.000 |
| Blautia | 0.036 | 0.026 | 8 | 6018 | 0.001 |
| Unknown | 0.036 | 0.020 | 17 | 6064 | 0.001 |
| Erysipelatoclostridium | 0.033 | 0.027 | 3489 | 7332 | 0.933 |
| Epulopiscium | 0.030 | 0.028 | 4027 | 4490 | 1.000 |
| Aeribacillus | 0.026 | 0.028 | 1148 | 5211 | 0.022 |
| Paenibacillus | 0.020 | 0.018 | 2340 | 5006 | 0.984 |
| Terrisporobacter | 0.013 | 0.014 | 649 | 2441 | 0.507 |
| Flavonifractor | 0.013 | 0.017 | 6 | 2153 | 0.001 |
| Enterobacter | 0.013 | 0.025 | 1 | 2247 | 0.001 |

*Table S12: Genera that contributed more than 1 per cent to the Bray-Curtis dissimilarity on day 5 between groups without and with antibiotic treatment in pens inoculated with ESBL. 'No' and 'Yes' refers to pens without and with antibiotics, respectively. Abbreviations: p: permutation p-value.*

| Genus | Average contribution | Standard deviation | Average abundance No | Average abundance Yes | p |
|---|---|---|---|---|---|
| Enterococcus | 0.142 | 0.093 | 1534 | 24727 | 0.001 |
| Bacillus | 0.081 | 0.105 | 70 | 12864 | 0.001 |
| Unknown | 0.079 | 0.070 | 7 | 12944 | 0.001 |
| Escherichia/Shigella | 0.071 | 0.048 | 17265 | 18512 | 1.000 |
| Blautia | 0.056 | 0.058 | 6 | 8685 | 0.001 |
| Clostridium sensu stricto 1 | 0.047 | 0.033 | 8585 | 9634 | 1.000 |
| Aeribacillus | 0.035 | 0.068 | 115 | 6019 | 0.001 |
| Klebsiella | 0.024 | 0.034 | 5 | 3999 | 0.001 |
| Erysipelatoclostridium | 0.018 | 0.014 | 4100 | 2683 | 1.000 |
| Paenibacillus | 0.017 | 0.012 | 4226 | 2864 | 1.000 |
| Epulopiscium | 0.014 | 0.021 | 1683 | 1314 | 0.999 |
| Flavonifractor | 0.014 | 0.011 | 317 | 2515 | 0.002 |
| Anaerostipes | 0.013 | 0.014 | 1927 | 1249 | 1.000 |
| Lachnoclostridium | 0.012 | 0.009 | 1 | 1902 | 0.001 |

*Table S13: Genera that contributed more than 1 per cent to the Bray-Curtis dissimilarity on day 5 between groups without and with antibiotic treatment in pens inoculated with catA1. 'No' and 'Yes' refers to pens without and with antibiotics, respectively. Abbreviations: p: permutation p-value.*

| Genus | Average contribution | Standard deviation | Average abundance No | Average abundance Yes | p |
|---|---|---|---|---|---|
| Clostridium sensu stricto 1 | 0.122 | 0.118 | 6410 | 24651 | 0.001 |
| Escherichia/Shigella | 0.068 | 0.056 | 16952 | 18135 | 1.000 |
| Bacillus | 0.064 | 0.067 | 58 | 11960 | 0.001 |
| Enterococcus | 0.062 | 0.071 | 9330 | 20546 | 0.588 |
| Blautia | 0.059 | 0.066 | 12 | 13291 | 0.001 |
| Unknown | 0.046 | 0.045 | 12 | 9629 | 0.001 |
| Unknown | 0.023 | 0.045 | 9 | 3900 | 0.001 |
| Epulopiscium | 0.022 | 0.020 | 4102 | 2349 | 1.000 |
| Thermoactinomyces | 0.021 | 0.058 | 0 | 3016 | 0.001 |
| Aeribacillus | 0.021 | 0.014 | 824 | 4300 | 0.003 |
| Lachnoclostridium | 0.018 | 0.028 | 4 | 3408 | 0.001 |
| Klebsiella | 0.014 | 0.029 | 3 | 2068 | 0.001 |
| Paenibacillus | 0.013 | 0.011 | 2255 | 2436 | 1.000 |

### 3. Background to the models

#### 3.1. Susceptible-infectious model

Environmental bacteria are the source of transmission in the susceptible-infectious model. The overall compartmental model is the following: susceptible animals ($S_t$) are infected by environmental bacteria ($B_t$) at a rate given by transmission rate parameter $\beta$:

$$\frac{dS_t}{dt} = -\beta S_t B_t \tag{1}$$

$$\frac{dI_t}{dt} = \beta S_t B_t \tag{2}$$

The environmental bacteria are produced by excreting animals ($I_t$) excreting viable bacteria into the environment at a constant rate of $\omega$ units per hour, and decay at rate $\delta$ per hour. As a result, the change in the amount of environmental bacteria at time t is given by:

$$\frac{dB_t}{dt} = \omega I_t - \delta B_t \tag{3}$$

This means that the number of bacteria in the environment at time *t* produced by an individual broiler *k* is defined by the time since the start of excretion of broiler *k*, $\tau_k$. The amounts of excreted bacteria are summed within each pen, such that the total amount of environmental bacteria at each time point *t* in a pen with *n* individuals is given by:

$$B_t = \sum_{k=1}^{n} \frac{\omega}{\delta}\left(1 - e^{-\delta\tau_k}\right) \tag{4}$$

The excretion rate ($\omega$) is unknown and unidentifiable in the inference method and therefore we set the excretion rate ($\omega$) to $\frac{\delta^2}{\delta + e^{-\delta} - 1}$ so that the total hazard produced by 1 excreting broiler during 1 time unit equals 1 (Gerhards et al., 2022). Thus, the hazard of colonization by environmental bacteria produced by all excreting individuals until that time, denoted $E_t$, is given by:

$$E_t = \frac{\delta^2}{\delta + e^{-\delta} - 1} B_t = \sum_{k=1}^{n} \frac{\delta}{\delta + e^{-\delta} - 1}\left(1 - e^{-\delta \cdot \tau_k}\right) \tag{5}$$

$\beta$ can be interpreted as the rate of transmission per time unit given the hazard $E_t = 1$.

#### 3.2. Bayesian hierarchical inference

We defined the likelihood function for the transmission rate parameter of each pen, with the number of new cases ($C_{i,t}$) binomially distributed with the probability of colonization ($p_i$) given the environmental hazard, where *i* is the numerical index for each pen.

$$C_i \sim Binomial(S_i, p_i) \tag{6}$$

The transmission rate parameter of each pen ($\beta$) was inferred with non-centred parameterization where the average transmission rate parameter of all pens ($\bar{a}$) is calculated and combined with the variation in the transmission rate parameter between pens ($z_i$).

$$\beta_i = \bar{a} + z_i \tag{7}$$

Posterior distributions of the transmission rate parameter for the different clusters (i.e., inoculum and antibiotic treatment) were obtained by combining the posterior distributions of $\bar{a} + z_i$ of all pens in that specific cluster.

This leads to the number of cases following a likelihood function (Eq. 6) combining the probability of transmission during the time interval between sampling points, $p_i\ (t \rightarrow t + \Delta t)$, environmental hazard ($E_t$), the decay rate ($\delta$) and input data. In the likelihood function $E_t\left(1 - e^{-\delta\Delta t}\right)$ characterizes a cumulative exposure during the sample interval by the amount of viable environmental bacteria at the beginning of the sampling interval and $I_t \frac{\delta^2}{\delta + e^{-\delta} - 1}\left(\frac{e^{-\delta\Delta t} - 1}{\delta} + \Delta t\right)$ is the cumulative exposure of individuals ($I_t$) by environmental bacteria during the sampling interval. $I_t$ are all inoculated and contact broilers excreting resistant bacteria during a time interval and all animals were assumed to have the same level of infectivity.

$$E(C_t + \Delta t) = S_t \times p(t \rightarrow t + \Delta t) = S_t\left(1 - e^{-\frac{\beta}{\delta}\left(E_t\left(1 - e^{-\delta\Delta t}\right) + I_t \frac{\delta^2}{\delta + e^{-\delta} - 1}\left(\frac{e^{-\delta\Delta t} - 1}{\delta} + \Delta t\right)\right)}\right) \quad (6)$$

##### 3.3. Decay rate

Broilers were excreting until the end of the experiment (Table S1), making it impossible to estimate the decay rate of *E. coli* by sampling from a prior probability distribution and simultaneously estimate the transmission rate with the Bayesian model in our study because a given number of cases can be explained equally well by a higher transmission rate or by a lower decay rate. We reviewed the literature on decay rates (Table S14) to find a suitable range of decay rates and ran the hierarchical model with several fixed decay rates ranging from 0.04 – 55 $h^{-1}$ (Table S15). This entire range of decay rates could be fitted well with low Watanabe–Akaike information criterion and divergence transition. Multiple studies in various environments suggest a very low level of *E. coli* decay in the first few days (Burrows and Rankin, 1970; Kovács and Tamási, 1977; Rogers et al., 2011), and *E. coli* excreted from broilers was found to have a decay rate of zero for 14 days (van Bunnik et al., 2014). A decay rate of zero could not be used with the model, therefore we selected the lowest fixed decay rate ($\delta$) of 0.04 $h^{-1}$ in the final model.

*Table S14: Decay of Escherichia coli outside a live host. Abbreviations: fc: field capacity.*

| ID | Temperature (°C) | Humidity | pH | Environment | Decay rate (per day) | Reference |
|---|---|---|---|---|---|---|
| 1 | Autumn | | | Lab | 0.102 | (Burrows and Rankin, 1970) |
| 2 | Autumn | | | Lab | 0.287 | (Burrows and Rankin, 1970) |
| 3 | 4 | | 7 | Lab | 0.686 | (Kovács and Tamási, 1977) |
| 4 | January | | | Lab | 0.109 | (Rankin and Taylor, 1969) |
| 5 | 26 | | 7.4 | Soil | 0.896 | (Klein and Casida, 1967) |
| 6 | 10 | | 7.4 | Soil | 0.195 | (Klein and Casida, 1967) |
| 7 | | | | Soil | 0.115 | (Mallmann and Litsky, 1951) |
| 8 | | | 7 | Soil | 0.371 | (Van Donsel et al., 1967) |
| 9 | | | | Soil | 0.143 | (Mallmann and Litsky, 1951) |
| 10 | | 1/3 bar | 6.16 | Soil | 0.473 | (Tate, 1978) |
| 11 | | saturated | 6.64 | Soil | 0.839 | (Tate, 1978) |
| 12 | | 100% fc | 6.16 | Soil | 0.796 | (Tate, 1978) |
| 13 | | flooded | | Soil | 0.382 | (Tate, 1978) |
| 14 | 0 | | | Inoculated water | 0.192 | (Mitchell, 1968) |
| 15 | 10 | 60% fc | | Swine manure-amended soil | 0.22 | (Rogers et al., 2011) |
| 16 | 10 | 80% fc | | Swine manure-amended soil | 0.19 | (Rogers et al., 2011) |
| 17 | 25 | 60% fc | | Swine manure-amended soil | 0.40 | (Rogers et al., 2011) |
| 18 | 25 | 80% fc | | Swine manure-amended soil | 0.28 | (Rogers et al., 2011) |
| 19 | 10 | 60% fc | | Beef manure-amended soil | 0.17 | (Rogers et al., 2011) |
| 20 | 10 | 80% fc | | Beef manure-amended soil | 0.15 | (Rogers et al., 2011) |
| 21 | 25 | 60% fc | | Beef manure-amended soil | 0.33 | (Rogers et al., 2011) |
| 22 | 25 | 80% fc | | Beef manure-amended soil | 0.37 | (Rogers et al., 2011) |
| 23 | Optimal condition to rear broilers | | | Broiler pen floor | 0.0 | (van Bunnik et al., 2014) |

*Table S15: Transmission rate parameters per hour for different decay rates of Escherichia coli outside a live host.*

| Decay rate ($h^{-1}$) | CPE without amoxicillin | ESBL without amoxicillin | catA1 without amoxicillin | CPE with amoxicillin | ESBL with amoxicillin | catA1 with amoxicillin |
|---|---|---|---|---|---|---|
| 0.04 | 0.0001 [0.0001, 0.0003] | 0.0003 [0.0001, 0.0005] | 0.0003 [0.0002, 0.0005] | 0.0004 [0.0002, 0.0008] | 0.0010 [0.0005, 0.0024] | 0.0008 [0.0004, 0.0024] |
| 0.05 | 0.0002 [0.0001, 0.0003] | 0.0003 [0.0002, 0.0006] | 0.0003 [0.0002, 0.0006] | 0.0005 [0.0002, 0.0009] | 0.0012 [0.0006, 0.0026] | 0.0009 [0.0005, 0.0026] |
| 0.14 | 0.0004 [0.0003, 0.0008] | 0.0006 [0.0004, 0.0013] | 0.0007 [0.0004, 0.0013] | 0.0010 [0.0005, 0.0019] | 0.0021 [0.0011, 0.0048] | 0.0018 [0.0010, 0.0052] |
| 0.17 | 0.0006 [0.0003, 0.0011] | 0.0008 [0.0005, 0.0015] | 0.0008 [0.0005, 0.0015] | 0.0012 [0.0007, 0.0023] | 0.0024 [0.0013, 0.0057] | 0.0021 [0.0012, 0.0061] |
| 0.22 | 0.0008 [0.0004, 0.0015] | 0.0011 [0.0006, 0.0021] | 0.0012 [0.0006, 0.0021] | 0.0017 [0.0009, 0.0031] | 0.0034 [0.0017, 0.0073] | 0.0028 [0.0016, 0.0086] |
| 0.27 | 0.0010 [0.0006, 0.0018] | 0.0014 [0.0008, 0.0026] | 0.0014 [0.0008, 0.0025] | 0.0021 [0.0011, 0.0040] | 0.0041 [0.0021, 0.0088] | 0.0036 [0.0020, 0.0104] |
| 0.37 | 0.0014 [0.0008, 0.0026] | 0.0018 [0.0011, 0.0037] | 0.0018 [0.0011, 0.0036] | 0.0028 [0.0016, 0.0054] | 0.0057 [0.0029, 0.0120] | 0.0051 [0.0027, 0.0141] |
| 0.45 | 0.0018 [0.0010, 0.0032] | 0.0023 [0.0013, 0.0045] | 0.0024 [0.0014, 0.0044] | 0.0034 [0.0019, 0.0068] | 0.0063 [0.0036, 0.0147] | 0.0063 [0.0034, 0.0172] |
| 0.61 | 0.0025 [0.0013, 0.0045] | 0.0033 [0.0018, 0.0063] | 0.0033 [0.0019, 0.0061] | 0.0048 [0.0027, 0.0093] | 0.0086 [0.0049, 0.0200] | 0.0087 [0.0047, 0.0246] |
| 0.74 | 0.0030 [0.0017, 0.0056] | 0.0041 [0.0023, 0.0077] | 0.0042 [0.0024, 0.0075] | 0.0061 [0.0033, 0.0117] | 0.0113 [0.0062, 0.0251] | 0.0109 [0.0056, 0.0308] |
| 1.00 | 0.0043 [0.0024, 0.0076] | 0.0056 [0.0032, 0.0108] | 0.0055 [0.0033, 0.0104] | 0.0079 [0.0046, 0.0159] | 0.0157 [0.0084, 0.0344] | 0.0146 [0.0080, 0.0422] |
| 1.35 | 0.0056 [0.0032, 0.0105] | 0.0076 [0.0043, 0.0147] | 0.0071 [0.0044, 0.0143] | 0.0114 [0.0062, 0.0214] | 0.0209 [0.0112, 0.0480] | 0.0193 [0.0111, 0.0567] |
| 1.65 | 0.0071 [0.0039, 0.0131] | 0.0093 [0.0052, 0.0183] | 0.0097 [0.0054, 0.0175] | 0.0145 [0.0076, 0.0269] | 0.0252 [0.0141, 0.0576] | 0.0243 [0.0130, 0.0679] |
| 2.23 | 0.0090 [0.0054, 0.0178] | 0.0124 [0.0071, 0.0247] | 0.0131 [0.0074, 0.0240] | 0.0181 [0.0104, 0.0362] | 0.0353 [0.0190, 0.0771] | 0.0325 [0.0182, 0.0944] |
| 2.72 | 0.0118 [0.0066, 0.0216] | 0.0162 [0.0088, 0.0305] | 0.0163 [0.0090, 0.0293] | 0.0229 [0.0131, 0.0447] | 0.0436 [0.0234, 0.0960] | 0.0411 [0.0220, 0.1140] |
| 3.67 | 0.0156 [0.0089, 0.0292] | 0.0223 [0.0120, 0.0410] | 0.0228 [0.0122, 0.0398] | 0.0310 [0.0173, 0.0602] | 0.0620 [0.0327, 0.1316] | 0.0572 [0.0298, 0.1572] |
| 4.48 | 0.0190 [0.0108, 0.0356] | 0.0267 [0.0143, 0.0500] | 0.0250 [0.0153, 0.0486] | 0.0371 [0.0210, 0.0736] | 0.0697 [0.0384, 0.1606] | 0.0698 [0.0357, 0.1978] |
| 6.05 | 0.0262 [0.0146, 0.0487] | 0.0360 [0.0200, 0.0683] | 0.0348 [0.0201, 0.0664] | 0.0527 [0.0288, 0.1000] | 0.0978 [0.0523, 0.2203] | 0.0885 [0.0500, 0.2644] |
| 7.39 | 0.0317 [0.0181, 0.0590] | 0.0446 [0.0237, 0.0838] | 0.0444 [0.0253, 0.0804] | 0.0616 [0.0352, 0.1213] | 0.1245 [0.0641, 0.2661] | 0.1168 [0.0626, 0.3162] |
| 20.09 | 0.0862 [0.0491, 0.1595] | 0.1250 [0.0649, 0.2245] | 0.1183 [0.0691, 0.2153] | 0.1739 [0.0938, 0.3321] | 0.3245 [0.1757, 0.7294] | 0.3069 [0.1697, 0.8615] |
| 54.60 | 0.2278 [0.1350, 0.4424] | 0.3181 [0.1797, 0.6136] | 0.3219 [0.1857, 0.5967] | 0.4699 [0.2545, 0.8954] | 0.8932 [0.4778, 1.9827] | 0.9289 [0.4530, 2.3571] |

##### 3.4. Posterior distribution of the model parameters

The posterior distribution is a multiplicative product of the prior distribution and the likelihood of producing the observed data. Figure S10 shows overlapping prior and posterior distribution for the average transmission rate parameter over all pens ($\bar{a}$; left) and the between-pen variation of the transmission rate parameter ($z_i$; right).

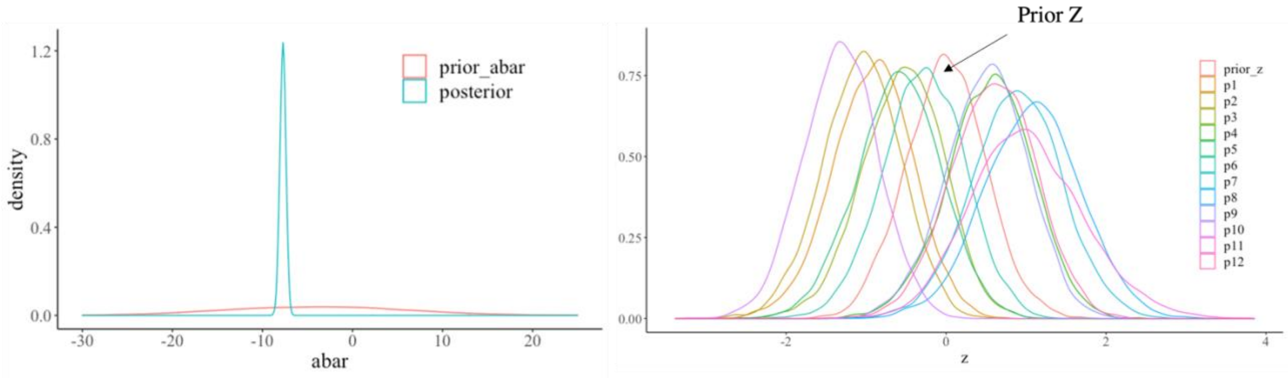


*Figure S10: Prior and posterior distributions for the average transmission rate parameter of all pens ($\bar{a}$; left) and the variation in the transmission rate parameters between pens ($z_i$; right) that are needed to obtain the transmission rate parameters ($\beta$).*

##### 3.5. Model diagnostics

Four thousand samples were drawn from each chain in which the first 2000 samples were warm-up samples. The model resulted in effective sample sizes (*ESS*) of 1200 in the average transmission rate over all pens ($\bar{a}$) and 2200 in the between-pen variation of the transmission rate ($z_i\sigma$). This effective sample size is above the number of samples, indicating efficient sampling of the posterior distribution. Gelman-Ruben convergences ($\hat{R}$) of 1 indicated that the Markov Chains converged. Similarly, patterns of the chains showed they rapidly explore a wide distribution and have a similar central tendency with a similar location with high probability, showing their good mixing, stability and convergence (McElreath, 2020; Stan Development Team) (Figure S11).

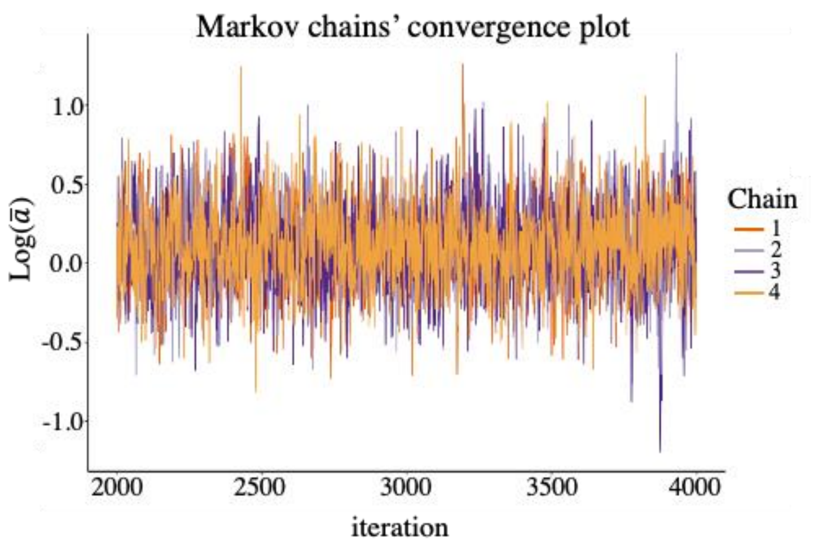


*Figure S11: Samples from each Markov chain are plotted sequentially in a trace plot. The horizontal axis is the serial number of draws (the first 2000 draws are the warm-up and have been removed) and the y-axis is the parameter value.*

#### 4. Plasmids and resistant genes

ResFinder 4.1 (Florensa et al., 2022) and PlasmidFinder 2.0 (Carattoli et al., 2014) were used to characterize the resistance genes and plasmids in the different inoculums. The CPE strain used in the animal experiment contained resistance genes from 6 families (*Table S16*) and 3 plasmids with incompatibility types IncHI2, IncHI2A, and IncQ1. The ESBL strain contained resistance genes from 4 families (*Table S17*) and 6 plasmids with incompatibility types ColpVC, IncFIB, IncFII, IncHI2, IncHI2A, and IncQ1. The catA1 strain contained resistance genes from 4 families (*Table S18*) and 4 plasmids with incompatibility types IncFIB, IncFII, IncI1-I, and IncI2.

*Table S16: The resistance genes present in the CPE strain.*

| Antimicrobial class | Genetic background | Antimicrobial resistance |
|---|---|---|
| amphenicol | floR | chloramphenicol |
| beta-lactam | blaOXA-162 | cefepime, ertapenem, imipenem, meropenem, piperacillin, tazobactam |
| beta-lactam | blaTEM-1B | ampicillin |
| folate pathway antagonist | sul2 | sulfamethoxazole |
| polymyxin | mcr-1.1 | colistin |
| quinolone | gyrA | ciprofloxacin, nalidixic acid |
| tetracycline | tet(A) | tetracycline |

*Table S17: Resistance genes present in the ESBL strain.*

| Antimicrobial class | Genetic background | Antimicrobial resistance |
|---|---|---|
| beta-lactam | blaCTX-M-2 | cefepime, cefotaxime, ceftazidime |
| beta-lactam | blaCTX-M-2, blaTEM-1B | ampicillin |
| folate pathway antagonist | dfrA1 | trimethoprim |
| folate pathway antagonist | sul1, sul2 | sulfamethoxazole |
| quinolone | gyrA | ciprofloxacin, nalidixic acid |
| tetracycline | tet(A), tet(B) | tetracycline |

*Table S18: Resistance genes present in the catA1 strain.*

| Antimicrobial class | Genetic background | Antimicrobial resistance |
|---|---|---|
| amphenicol | catA1 | chloramphenicol |
| beta-lactam | blaTEM-1B | ampicillin |
| folate pathway antagonist | dfrA1 | trimethoprim |
| | sul2 | sulfamethoxazole |
| tetracycline | tet(A) | tetracycline |

#### 5. References

Burrows, M.R., Rankin, J.D., 1970. A further examination of the survival of pathogenic bacteria in cattle slurry. Br Vet J 126, xxxii+.

Carattoli, A., Zankari, E., García-Fernández, A., Voldby Larsen, M., Lund, O., Villa, L., Møller Aarestrup, F., Hasman, H., 2014. In silico detection and typing of plasmids using PlasmidFinder and plasmid multilocus sequence typing. Antimicrob Agents Chemother 58, 3895-3903.

Florensa, A.F., Kaas, R.S., Clausen, P.T.L.C., Aytan-Aktug, D., Aarestrup, F.M., 2022. ResFinder – an open online resource for identification of antimicrobial resistance genes in next-generation sequencing data and prediction of phenotypes from genotypes. Microbial Genomics 8.

Gerhards, N.M., Gonzales, J.L., Vreman, S., Ravesloot, L., van den Brand, J.M.A., Doekes, H.P., Egberink, H.F., Stegeman, A., Oreshkova, N., van der Poel, W.H.M., de Jong, M.C.M., 2022. Efficient direct and limited environmental transmission of SARS-CoV-2 lineage B.1.22 in domestic cats. bioRxiv, 2022.2006.2017.496600.

Klein, D.A., Casida, L.E., Jr., 1967. Escherichia coli die-out from normal soil as related to nutrient availability and the indigenous microflora. Can J Microbiol 13, 1461-1470.

Kovács, F., Tamási, G., 1977. Survival times of bacteria in liquid manure. Acta Vet Acad Sci Hung 27, 47-54.

Mallmann, W.L., Litsky, W., 1951. Survival of selected enteric organisms in various types of soil. Am J Public Health Nations Health 41, 38-44.

McElreath, R., 2020. Statistical rethinking: a bayesian course with examples in R and Stan. Chapman & Hall.

Mitchell, R., 1968. Factors affecting the decline of non-marine micro-organisms in seawater. Water Research 2, 535-543.

Rankin, J.D., Taylor, R.J., 1969. A study of some disease hazards which could be associated with the system of applying cattle slurry to pasture. Vet Rec 85, 578-581.

Rogers, S.W., Donnelly, M., Peed, L., Kelty, C.A., Mondal, S., Zhong, Z., Shanks, O.C., 2011. Decay of bacterial pathogens, fecal indicators, and real-time quantitative PCR genetic markers in manure-amended soils. Applied and Environmental Microbiology 77, 4839-4848.

Stan Development Team, 2022. Runtime warnings and convergence problems.

Tate, R.L., 3rd, 1978. Cultural and environmental factors affecting the longevity of Escherichia coli in Histosols. Appl Environ Microbiol 35, 925-929.

van Bunnik, B.A., Ssematimba, A., Hagenaars, T.J., Nodelijk, G., Haverkate, M.R., Bonten, M.J., Hayden, M.K., Weinstein, R.A., Bootsma, M.C., De Jong, M.C., 2014. Small distances can keep bacteria at bay for days. Proc Natl Acad Sci U S A 111, 3556-3560.

Van Donsel, D.J., Geldreich, E.E., Clarke, N.A., 1967. Seasonal variations in survival of indicator bacteria in soil and their contribution to storm-water pollution. Appl Microbiol 15, 1362-1370.
